# Multiplexed proximity labelling proteomics identifies a non-canonical Bcl-2 family interactome associated with apoptotic priming

**DOI:** 10.1101/2025.05.30.657000

**Authors:** Robert Pedley, Charlotte E. L. Mellor, Louise E. King, Matthew Jones, Isobel Taylor-Hearn, Alexander Mironov, Craig Lawless, Andrew P. Gilmore

**Author notes:** Correspondence to: Dr. Andrew P. Gilmore, Wellcome Centre for Cell-Matrix Research, Faculty of Biology, Medicine and Health, A.3034 Michael Smith Building, University of Manchester, Oxford Road, Manchester M13 9PT. These authors contributed equally to this work.

## Abstract

Bcl-2 family proteins govern the intrinsic apoptotic pathway by regulating mitochondrial outer membrane permeabilisation (MOMP), releasing apoptogenic factors into the cytosol. How close a cell is to MOMP, termed mitochondrial priming, is determined by the interactions between different Bcl-2 proteins at the outer mitochondrial membrane (OMM). However, Bcl-2 proteins can also drive diverse processes that do not result in cell death, such as incomplete MOMP, sublethal caspase activation, pro-inflammatory signalling and regulation of cellular metabolism. To understand the wider Bcl-2 family interactome and how it might be involved in these processes, we undertook an unbiased proteomic BioID screen, using both pro- and anti-apoptotic Bcl-2 family members as bait proteins. We found that most high-confidence potential interactions in non-apoptotic cells were outside the canonical Bcl-2 family interactome. Analysis of how this interactome changed in response to either BH3-mimetics or full-length BH3-only proteins revealed dynamic changes at multiple organelles and membrane contact sites in response to altered mitochondrial priming. These findings underscore the complex and varied interactions of the Bcl-2 family proteins, expanding their scope and function beyond the frequently studies intra-family interactions.

## Introduction

The Bcl-2 family of proteins regulate the intrinsic pathway of apoptosis, the point of commitment for which is mitochondrial outer membrane permeabilization (MOMP). MOMP releases apoptogenic proteins from the intermembrane space, activating caspases and driving cell death ^1^. However, the canonical view of apoptosis as an all- or nothing response to a lethal stress has been challenged. Minority or incomplete MOMP (iMOMP), where a fraction of mitochondria within a cell release cytochrome *c,* engages sublethal caspase activation and the innate immune response, regulates cell functions without apoptosis occurring ^2–5^. Thus, Bcl-2 proteins can function on mitochondria without commitment to cell death.

Prior to MOMP, cells vary in their degree of apoptotic sensitivity and can rapidly fluctuate between being relatively sensitive or resistant to apoptosis. Furthermore, cells show significant inter- and intraline variation in their sensitivity to MOMP and apoptosis ^6,7^. These variations in sensitivity are linked to the dynamic interactions of Bcl-2 proteins with the outer mitochondrial membrane (OMM) and each other ^7–9^. Our canonical understanding of Bcl-2 family interactions and MOMP is largely based on subjecting cells to a lethal stress. Thus, pro-death signals activate the pro-apoptotic effector Bcl-2 proteins Bax and Bak, which form pores in the OMM, whilst simultaneously inhibiting anti-apoptotic family members like Bcl-2 and Bcl-XL. Anti- and pro-apoptotic Bcl-2 proteins are themselves regulated by the BH3-only proteins, primarily via docking of the α-helical BH3-domain of the BH3-only proteins into a groove on the surface of the effectors. This interaction forms the rationale for BH3-mimetic drugs like venetoclax ^10^. However, this canonical view of Bcl-2 proteins does not fully explain their role in non-apoptotic cells. For example, many Bcl-2 family proteins retrotranslocate from mitochondria to the cytosol, the regulation of which alters sensitivity to MOMP ^7,9^.

Alongside the established intrafamily Bcl-2 protein interactions, several non-canonical partners have been reported. These include elements of the mitochondrial fusion and fission machinery, such as Drp1 ^11^, and FK506-binding protein 8 (FKBP8), which promotes targeting of Bcl-2 and Bcl-XL to the OMM ^12^. Several non-canonical interactions with BH3-only proteins have been identified, including Bad with 14.3.3 protein isoforms and glucokinase ^13,14^, and Bid with VDAC2 and MTCH2 ^15,16^. Some of these interactions have roles in apoptosis, whilst others have been linked to non-apoptotic functions of Bcl-2 proteins. However, a holistic understanding of this wider Bcl-2 family interactome in non-apoptotic cells is lacking.

Several hurdles exist for interrogating native Bcl-2 protein interactions. Different family members shuttle between distinct subcellular localisations ^7,9^, with both cytosolic and membrane-bound states that may be differentially sensitive to specific isolation conditions ^17,18^. Live imaging approaches have provided insight into native interactions, but are restricted to already known binding partners ^19^. An alternative, unbiased approach to identifying potential interactions is proximity biotin labelling identification (BioID) ^20^. BioID utilises a mutant *E. coli* biotin ligase, BirA*, which generates reactive biotinoyl-AMP in the presence of excess biotin. Biotinoyl-AMP is short-lived and can covalently label lysine residues within an approximately 10 nm radius of the enzyme. When BirA* is expressed as a fusion with a bait protein, labelling favours direct binding partners or proximal components of multiprotein complexes of that bait (prey proteins). BioID has been used to define interactions not amenable to conventional isolation techniques, such as multi-protein membrane complexes ^21^. In particular, it has been successfully used to interrogate the mitochondrial proteome ^22^.

Here we employed BioID to investigate the Bcl-2 family interactome in live cells subjected to dynamic shifts in apoptotic priming. We found that most prey proteins identified in non-apoptotic cells did not fall within the canonical Bcl-2 interactome, although several have been previously reported in the literature. Bax, Bcl-XL and Bcl-2 showed distinct but overlapping high-confidence proximal interactions that reflected their subcellular distribution. In contrast, the BH3-only protein Bid identified a limited number of predominantly canonical binding partners. Interestingly, inducing apoptotic priming by addition of either the full-length BH3-only protein Bad or the Bad BH3-mimetic ABT-737 produced distinct patterns of enrichment for Bcl-XL and Bcl-2 proximity interactions which correlated with distinct changes in cellular pathways. These data reveal a complex and dynamic Bcl-2 family interactome which fluctuates according to the level and source of priming and highlights their broader functional role in non-apoptotic cells.

## Results

### BioID identifies an in-situ interaction network for Bid, Bax and Bcl-XL in unprimed cells

To obtain a holistic view of the Bcl-2 family interactome within non-apoptotic cells we employed BioID, which allows *in situ* conjugation of biotin to proteins proximal to a BirA*-bait fusion^20^. Labelled proteins are isolated by streptavidin affinity purification and quantified by liquid chromatography/tandem mass spectrometry (LC-MS/MS, **Fig.1A**). We chose three bait proteins for initial analysis, Bcl-XL, Bax and Bid, selected for overlap in their STRING-predicted interaction networks. Bcl-XL represented a multi-domain anti-apoptotic protein with a broad binding profile for other Bcl-2 family members. Bax was chosen as a pro-apoptotic effector with a dynamic subcellular distribution. Bid was selected as it is a constitutively expressed BH3-only protein which binds both pro- and anti-apoptotic multidomain family members. Each bait was cloned into a lentiviral expression vector, pCDH, to co-express the myc epitope tagged BirA* bait fusion and tagBFP (**Fig.1A**). tagBFP facilitated normalising expression of each stably expressed bait by FACS. Venus-BirA* was used as a negative control bait. To verify the functionality of the BirA* tagged baits, Bax/Bak double knockout mouse embryonic fibroblasts (DKO MEFs) were transiently transfected with mCherry-tBid alone or with either pVenus-Bax or pCHD-tagBFP-BirA*-Bax, and apoptosis quantified (**Fig.1B**). mCherry-tBid induced cell death equally when co-expressed with either Venus-Bax or BirA*-Bax. Conversely, expression of mCherry-tBid induced apoptosis in wild-type (WT) MEFs, which was inhibited equally by either Venus-Bcl-XL or BirA*-Bcl-XL.

**Figure 1.**
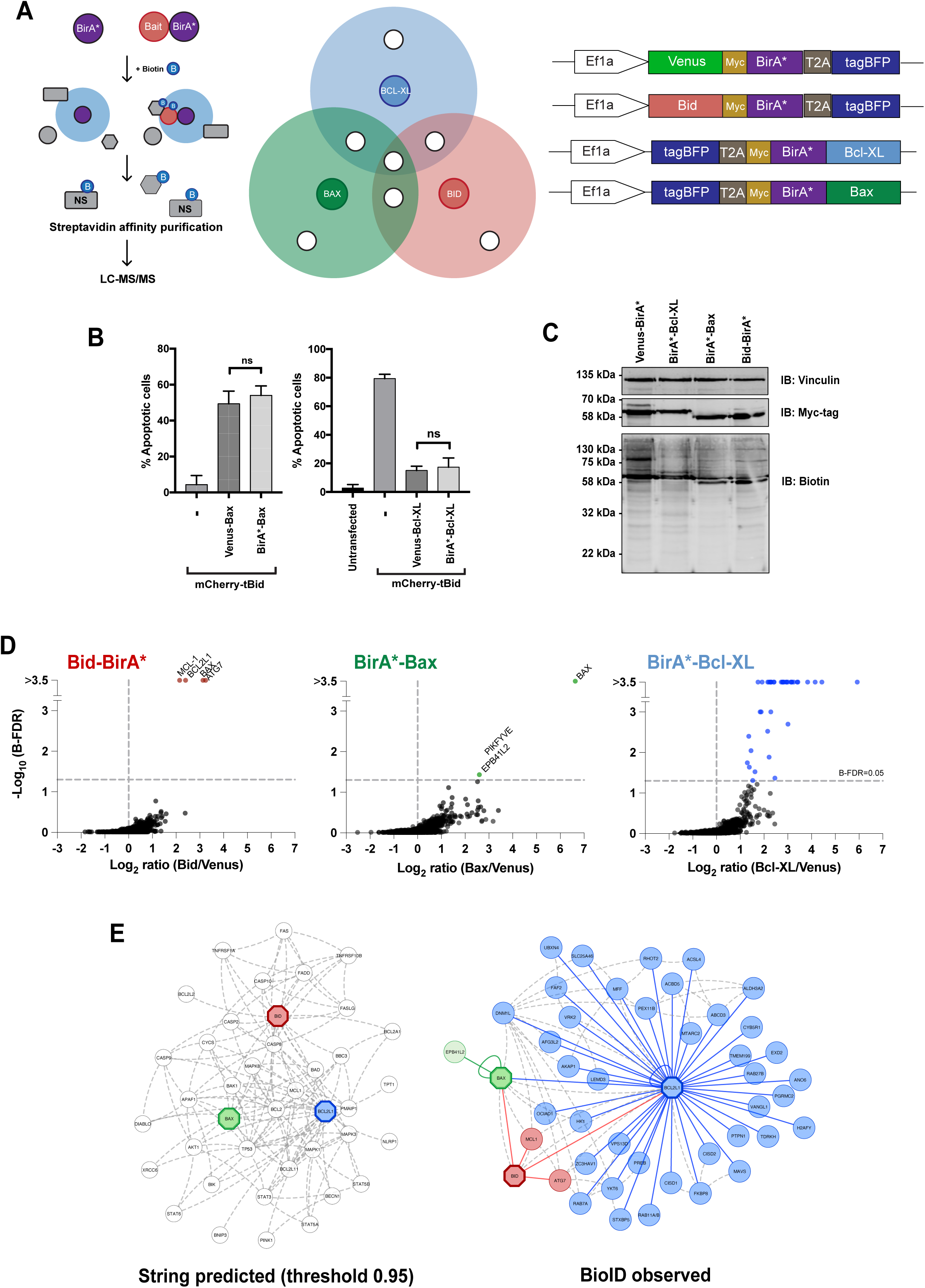
BioID identification of Bcl-2 protein family interactions in non-apoptotic cells. **A.** Schematic representation of the BioID workflow, with Venn diagram representation of expected prey overlap. Right panel shows schematic representation of BioID expression constructs. **B.** Functional validation of BirA*-Bax and BirA*-XL bait proteins. Left panel - Bax/Bak double knockout mouse embryonic fibroblasts (DKO MEFs) were transfected with mCherry-tBid either alone, or in combination with either Venus-Bax or BirA*-Bax. Right panel - wildtype MEFs were transfected with mCherry-tBid, either alone or in combination with Venus-Bcl-XL or BirA*-Bcl-XL. 18 hours post-transfection cells were fixed and apoptosis quantified. Data represent mean and SD of three independent experiments. Significance calculated by ANOVA, followed by Tukey’s multiple comparison test. Ns = non-significant. **C.** MCF10A cells stably expressing either Venus-BirA*, BirA*-Bcl-XL, BirA*-Bax, Bid-BirA* were incubated in 50 mM biotin for 18 hours. Whole-cell lysates were prepared and examined by immunoblotting for the indicated antibodies and streptavidin. **D.** Significance Analysis of Interactome (SAINT) identifies BioID preys significantly enriched in cells expressing Bid-BirA*, BirA*-Bax or BirA*-Bcl-XL, each relative to Venus-BirA* expressing cells. Note: for brevity, and to keep nomenclature consistent with later analyses, gene ID names are displayed when referring to prey proteins identified via BioID experiments/SAINT. **E.** Comparison of the predicted interactomes of Bid, Bax and Bcl-XL to the network observed through the BioID data. Left panel - network of predicted interactions generated using data derived from the STRING database. Proteins assigned a confidence score of 0.95 or higher via STRING are displayed. Right panel - network of genes identified via BioID experiments. Predicted interactions between genes within the observed network (STRING score of 0.4 or greater) are denoted as dashed grey lines.

Stable cell lines expressing each BioID bait or Venus-BirA* control at comparable levels were generated in the non-transformed mammary epithelial cell line, MCF10A (**Fig.1C**). Cells were allowed to reach 80% confluency, then incubated in media supplemented with 50 µM biotin for 18 hours. Cells were switched back to biotin-free media before lysis and biotinylated prey proteins isolated on streptavidin agarose. Three independent biological replicates were analysed by label-free, quantitative LC-MS/MS, and approximately two thousand individual proteins were identified within each bait sample. SAINTexpress was used to calculate the Bayesian false discovery rates (BFDR) and identify high-confidence interacting partners or proximal proteins, using Venus-BirA* to control for non-specific interactions. Labelled proteins with a BFDR of ≤0.05 were considered to represent a high confidence interaction/proximity event, and from here on are referred to as “preys” (**Fig.1D**). Of the three baits, BirA*-Bax identified the fewest potential interactions, with only itself and one other prey displaying a BFDR of <0.05. One other protein, the phosphoinositide kinase PIKfyve, fell just below the imposed BFDR threshold (BFDR=0.055). These data were consistent with Bax being predominantly monomeric and cytosolic in unprimed cells. Bid-BirA* identified four potential interaction partners with BFDRs <0.05. Three of these, Bax, Bcl-XL and Mcl-1, were in Bid’s STRING-predicted interaction network (**Fig.1E**). The fourth was the autophagy regulator ATG7 ^23^. In marked contrast to Bax and Bid, BirA*-Bcl-XL identified a complex set of potential interactions, with around 40 prey proteins reaching the set BFDR threshold. These were predominantly membrane-associated proteins, although, aside from Bax and itself, none were within the STRING-predicted network for Bcl-XL. Although Bcl-XL retrotranslocates like Bax, its equilibrium favours the membrane associated state, possibly contributing to the increased number of preys reaching the BFDR threshold.

We assembled the three BioID prey lists into a putative interaction network and compared this with the STRING-predicted network (**Fig.1E**). Within the observed network, only Bax, Bcl-XL and Mcl-1 were shared with the predicted network. However, many components within the predicted network are linked to pro-apoptotic signalling, such as caspase 8 and components of the Fas signalling platform for Bid. In contrast, our BioID network was determined in live cells not exposed to a dominant, pro-apoptotic signal. These data indicate that the STRING-predicted interactions, and thus the assumed canonical interaction network of Bcl-2 proteins generally, may not reflect those that predominate in unprimed and non-apoptotic cells.

### Bcl-2 shows a distinct but overlapping interactome to that of Bcl-XL which reflects its subcellular distribution

Different Bcl-2 proteins localise to multiple organelles, such as Bcl-2 to the endoplasmic reticulum and nuclear membrane ^24^. Given these differences in organelle localisations of Bcl-2 family proteins, we asked if the proximity interactomes of Bcl-2 and Bcl-XL overlapped at mitochondria or highlighted distinct interaction networks (**Fig.2A**). MCF10A cells were generated expressing BirA*-Bcl-2 (**Fig.S2A-C**), labelled, and biotinylated proteins identified by LC-MS/MS (**Fig.2B**). BirA*-Bcl-2 identified a range of prey proteins that reflected its broader subcellular distribution compared to Bcl-XL (**Fig.2C**). To identify overlap of the Bcl-2 and Bcl-XL interactomes, we compared average intensity and fold change between significantly enriched prey proteins identified by Bcl-XL and Bcl-2 using hierarchical cluster analysis (**Fig.2D**). Self-labelling represented the highest signal for each bait. Comparing the average intensity highlighted a wider range of potential interactions for Bcl-2 than Bcl-XL, but with notable overlap between enriched preys (**Fig.2C**, **Fig.S2D**). However, subsets of preys were significantly more enriched by the different baits. Of note, several OMM associated proteins were more prominent with BirA*-Bcl-XL, including AKAP1, MAVS, the mitochondrial myosin Myo19, Bid, and the mitochondrial fission regulators Drp1 and MFF. The most enriched prey proteins for BirA*-Bcl2 included the endoplasmic reticulum chloride channel CLCC1, syntaxin 5, the ankyrin-LEM domain-containing protein Ankle2, which localises to both the ER and the nuclear envelope, and several nuclear envelope components (BANF1, emerin, TMEM201, SUN1 and lamin B). Interestingly, the only canonical prey identified by BirA*-Bcl2, apart from itself, was Bcl-XL. These differences between Bcl-2 and Bcl-XL were highlighted by gene ontology (GO) analysis (**Fig.2E, Fig.S2E,F**). For Bcl-2, there was significant overrepresentation of terms associated with endoplasmic reticulum, membrane dynamics and GTPases. In contrast, Bcl-XL identified terms associated more with mitochondria, apoptosis and membrane permeability.

**Figure 2.**
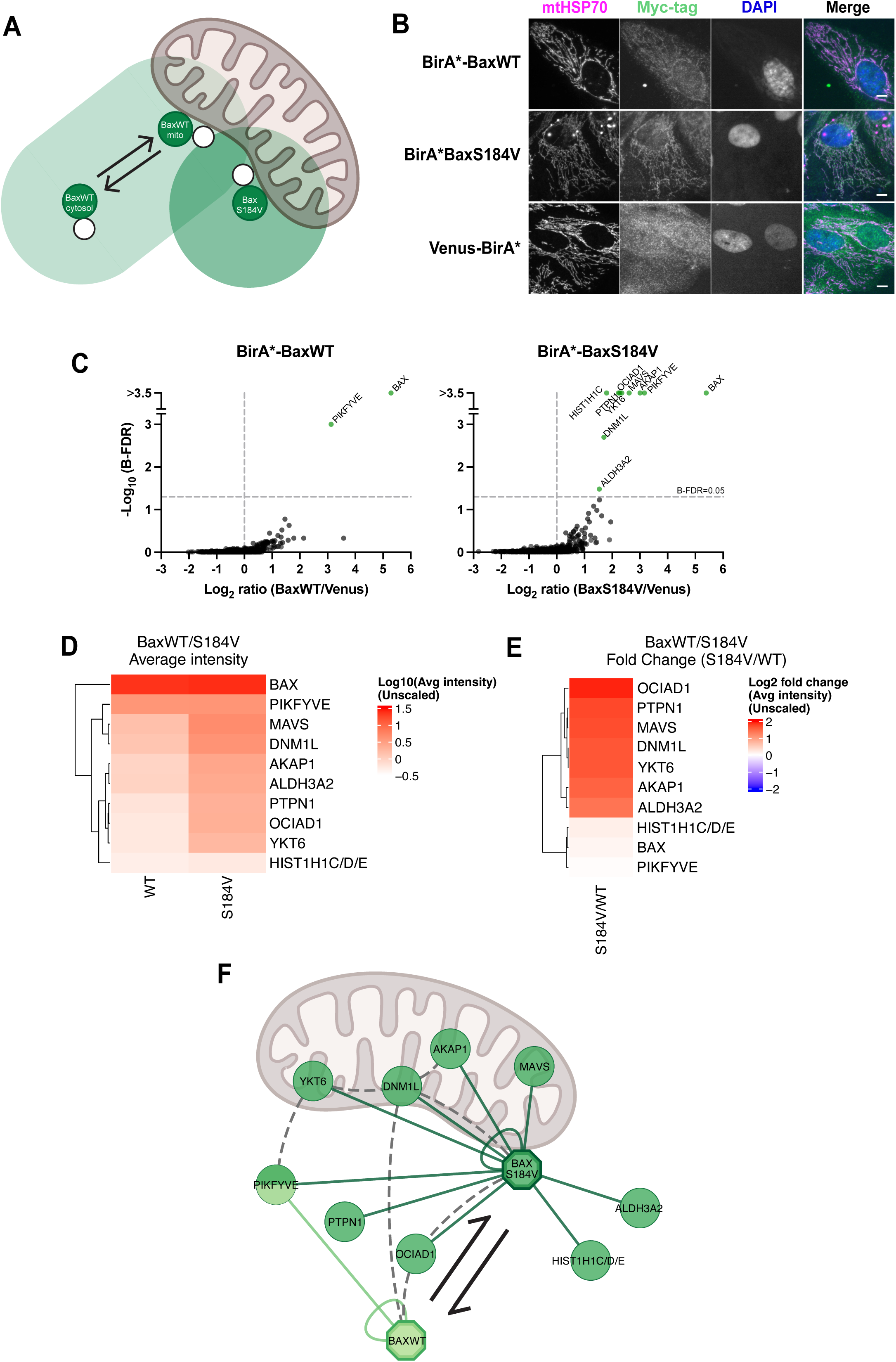
Proximity labelling indicates Bcl-2 has a wider subcellular distribution than Bcl-XL. **A.** Schematic representation of predicted BirA*-Bcl-2 and BirA*-Bcl-XL subcellular localisation and predicted labelling patterns. **B.** SAINT analysis identifies genes labelled in cells expressing either BirA*-Bcl-XL or BirA*-Bcl-2, relative to Venus-BirA* expressing cells. **C.** Network representation of Bcl-2 and Bcl-XL proximity interactomes. Preys identified as significantly enriched (B-FDR >0.05) by either BirA*-Bcl-2 or BirA*-Bcl-XL expressing cells in 3B are displayed. Preys significantly enriched by both baits are indicated. **D.** Heatmaps showing the average intensity of BioID presy identified by BirA*-Bcl-XL and Bcl-2 (left panel) and a comparison of relative enrichment of each prey by Bcl-2 compared to Bcl-XL (right panel). Preys are displayed if they reached significance (B-FDR > 0.05) for either BirA*-Bcl-2 or BirA*-Bcl-XL. Data represent the average of three independent experiments and are unscaled. **E.** Gene ontology analysis using Cellular Component annotations indicating the organelles most associated with the BioID preys identified by Bcl-2 and Bcl-XL.

These data indicated a broad subcellular distribution of Bcl-2 in unprimed epithelial cells, likely beyond that of a protein serving an exclusively anti-apoptotic function.

### Reduced Bax retrotranslocation identifies a mitochondrial interaction network overlapping with Bcl-XL

Bcl-XL, Bcl-2 and Bax are multidomain Bcl-2 family proteins that localise to membranes via a C-terminal tail anchor. However, BirA*-Bax identified few potential interactions in unprimed MCF10A cells. In unprimed cells mitochondrial Bax constitutively retrotranslocates from the OMM to the cytosol, where it is monomeric, potentially explaining the lack of prey proteins ^9^. As cells become primed through loss of survival signals, Bax retrotranslocation is inhibited and it accumulates on the OMM^9^. We therefore hypothesised that reduced Bax retrotranslocation would increase the number of identified prey proteins. To mimic Bax mitochondrial accumulation associated with priming we substituted serine 184 in the C-terminal tail anchor to valine (S184V), which reduces Bax retrotranslocation (**Fig.3A, S3A**) ^9^. We generated stable MCF10A cell lines expressing either BirA*-BaxWT or BirA*BaxS184V, Venus-BirA* was used as a control, and tagBFP to normalise expression. Immunofluorescence microscopy for the myc-tag verified expression and increased mitochondrial localisation of BirA*-BaxS184V compared to BirA*-BaxWT (**Fig.3B**). Biotin labelling and isolation of biotinylated proteins were performed as before (**Fig.S3B**). Preys enriched by either BirA*-BaxWT or BirA*-BaxS184V relative to Venus-BirA* were then identified and quantified by LC-MS/MS (**Fig.3C**). As before, few preys were enriched by BirA*-BaxWT, with itself again the predominant protein identified. PIKfyve was identified below the BFDR threshold of 0.05, consistent with the low BFDR value observed in the prior BioID experiment. In contrast, BirA*-BaxS184V identified ten preys, including itself and PIKyve.

**Figure 3.**
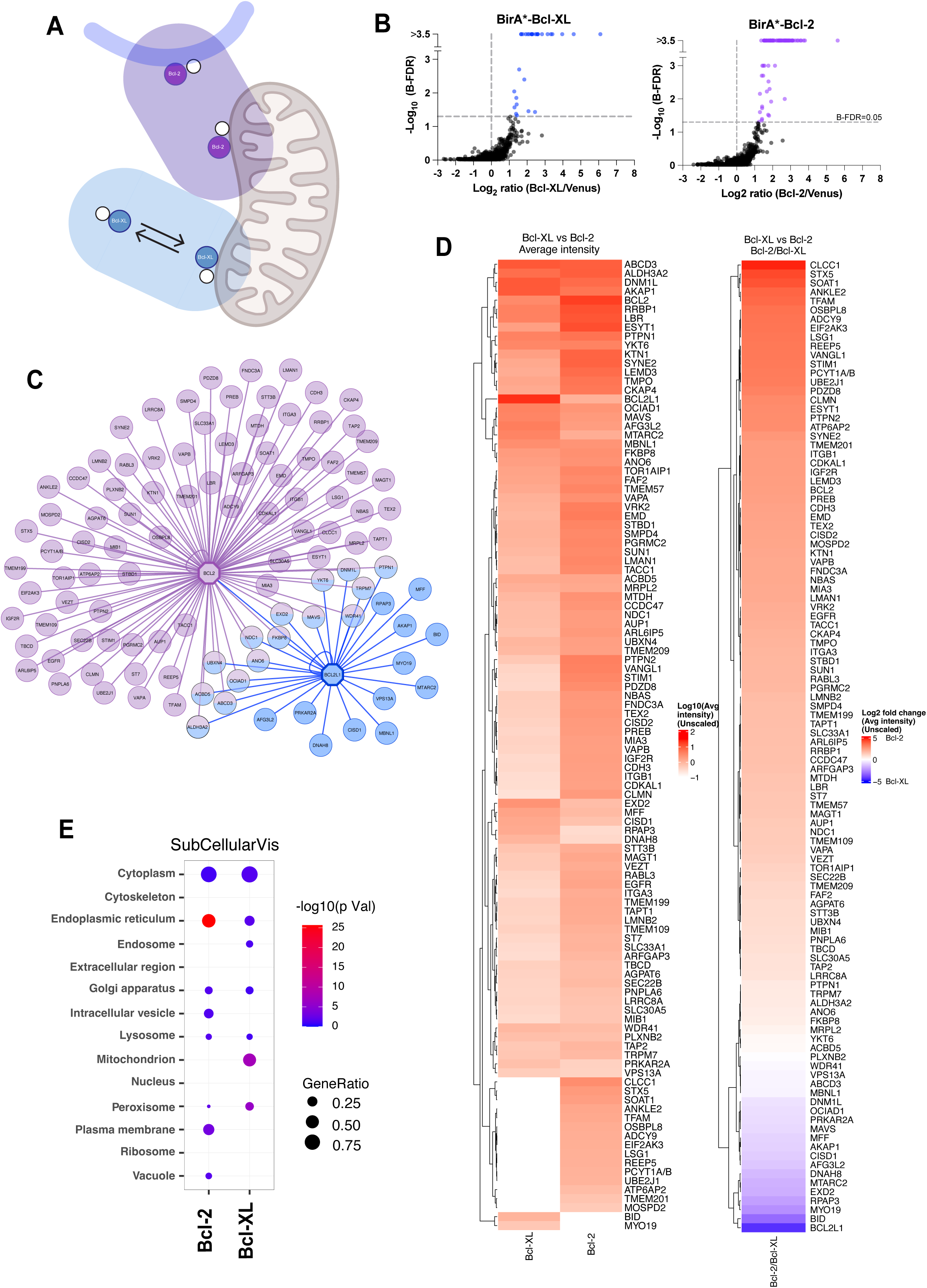
Bax retrotranslocation dependent mitochondrial interaction. **A.** Schematic representation of BirA*-BaxWT and BirA*-BaxS184V subcellular localisation, retrotranslocation dynamics and predicted labelling patterns. **B.** Representative images of MCF10A cells stably expressing either BirA*-BaxWT, BirA*-BaxS184V or Venus-BirA*. Cells were fixed and immunostained for the indicated proteins (mtHsp70 and Myc-tag). Scale bar represents 10 mm. **C.** BioID SAINT analysis identifies prey proteins significantly enriched by BirA*-BaxWT or BirA*-BaxS184V, relative to Venus-BirA* expressing cells. **D.** Heatmap representation of the average protein intensity for significant prey proteins. The average protein intensity for preys that had a SAINT derived BFDR of <0.05 (2C) are displayed for BirA*-BaxWT and BirA*-BaxS184V conditions. Data represent the mean of three independent experiments and are unscaled. **E.** Heatmap representation of the fold change of significant preys in Fig2C in BirA*-BaxS184V expressing cells relative to BirA*-BaxWT. Data source as described in Fig2D. **F.** Schematic representation of the prey protein network identified through BaxWT and BaxS184V proximity labelling. YKT6, DNM1L, AKAP1 and MAVS have shown strong previous evidence of mitochondrial localisation.

All of the prey proteins identified by BirA*-BaxS184V were present, albeit at lower abundance and with a BFDR >0.05, in the BirA*-BaxWT data set. We therefore compared both the average LC-MS/MS signal intensity and fold change relative to Venus-BirA* of these prey proteins between the two baits (**Fig.3D,E).** Comparing BirA*-BaxWT with BirA*-BaxS184V, there was no further enrichment of either Bax itself or PIKfyve when the bait was mitochondrial. However, several prey proteins were significantly enriched when Bax retrotranslocation was reduced. These included MAVS, DNM1L (Drp1), OCIAD1, YKT6, PTPN1 and AKAP1. We compared the STRING-predicted interactions of the enriched prey proteins identified by BirA*-BaxWT and BirA*-BaxS184V (**Fig.3F**). Some of the preys had predicted interactions with Bax, including Drp1. Other prey proteins were linked through STRING-curated interactions with each other, such as YKT6 and PIKfyve. Furthermore, we noted that many of the preys enriched by BirA*-BaxS184V were also identified by BirA*-Bcl-XL, including AKAP1, Drp1, MAVS, OCIAD1, PTPN1 and YKT6. Overall, these data suggested a potential core mitochondrial interactome around pro- and anti-apoptotic Bcl-2 family proteins that dynamically alters in response to protein retrotranslocation.

### Differential effects on the Bcl-2 family interactome when priming occurs through either full-length Bad or the Bad-mimetic ABT-737

We next asked how the Bcl-XL interactome might be influenced by shifts in mitochondrial priming. We recently described a conditionally active variant of full-length (FL) Bad, BadER, which can be post-translationally activated by the oestrogen analogue 4-hydroxytamoxifen (4-OHT) ^7^. In the presence of 4-OHT, BadER formed a stable complex with Bcl-XL on the OMM which correlated with increased apoptotic priming. BH3-mimetics are small molecule drugs designed to function in an analogous manner to BH3-only proteins, binding the same hydrophobic groove on anti-apoptotic Bcl-2 proteins and inhibiting their function. However, full-length BH3-only proteins can interact with multidomain Bcl-2 proteins in a manner distinct to BH3-mimetics ^25,26^. We therefore compared the effects of BadER and the BH3-mimetic, ABT-737, on the Bcl-XL interactome **(Fig.4A).** ABT-737 was chosen as it has a similar binding profile to the Bad BH3-domain, inhibiting Bcl-XL, Bcl-2 and Bcl-w ^10^.

**Figure 4.**
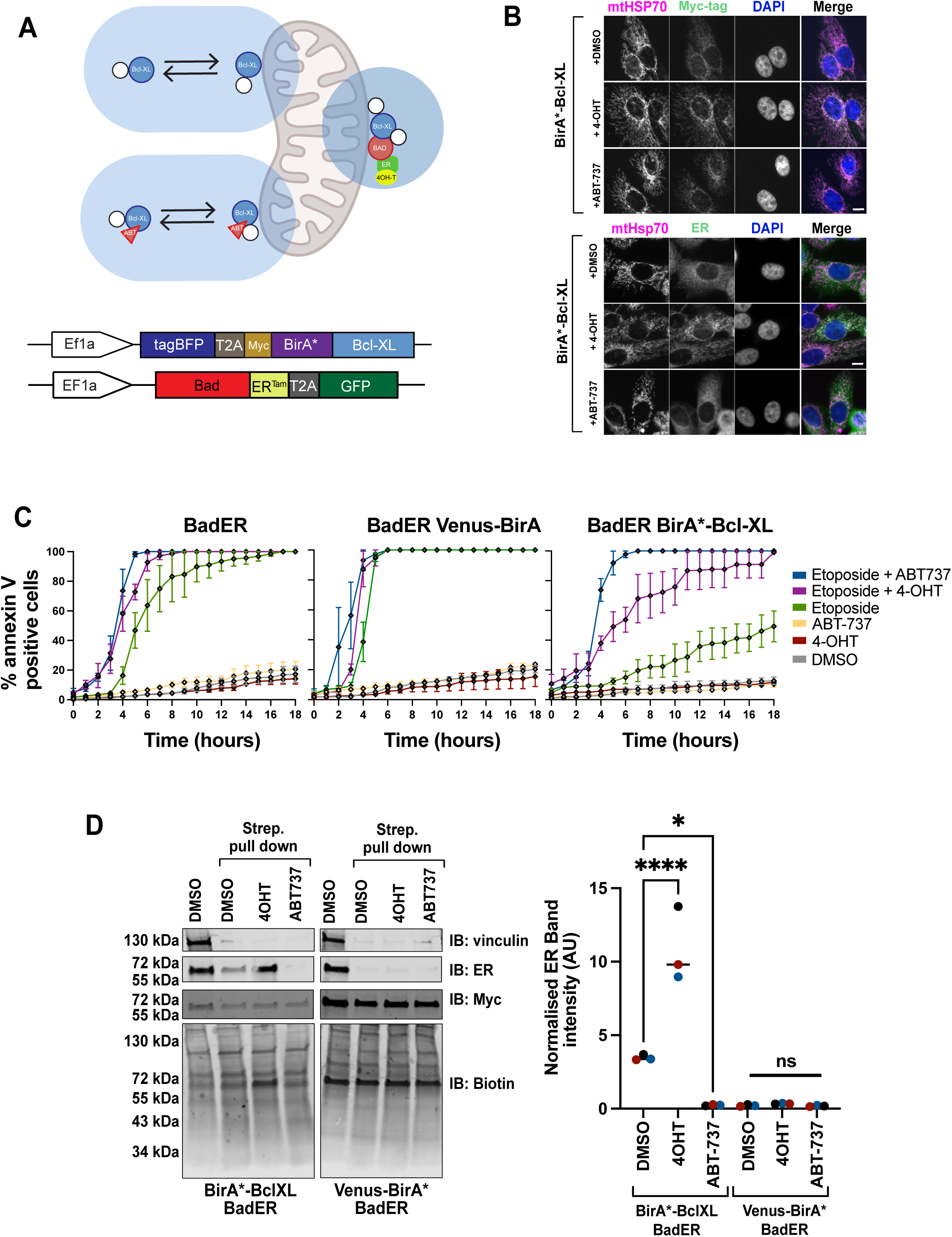
Generation of BirA*-Bcl-XL stable cells that can be induced into a primed state by a conditionally active BH3-only protein, BadER. **A.** Schematic representation of predicted BirA*-Bcl-XL subcellular localisation and predicted labelling patterns with 4-OHT and ABT-737 induced priming. Lower panel shows the BirA*-BclXL construct and the BadER-T2A-GFP lentiviral construct used to generate the MCF10A inducible priming lines, based upon published work ^7^. **B.** Immunofluorescence staining of MCF10A cells stably expressing BirA*-Bcl-XL and BadER, and treated with either DMSO, 10µM 4-OHT or 5µM ABT-737. In the top panel, cells were co-immunostained with anti-mtHSP70 and anti-myc tag, along with DAPI. Bottom panel - cells were co-immunostained for the anti-oestrogen receptor (ER) tag to show the BadER fusion protein, and anti-mtHSP70, in addition to DAPI. Scale bar = 10µm. **C.** MCF10A cells expressing BadER, with or without either Venus-BirA* or BirA*-Bcl-XL, were treated with DMSO, 10 µM 4-OHT, or 5 µM ABT-737, with or without 800 µM etoposide. Cells were imaged hourly over 18 hours in the presence of FITC Annexin V and PI, and cumulative Annexin V positivity was calculated as a percentage of the initial cell population. Data represent the mean ± SEM of three independent experiments. **D.** MCF10A cells expressing BadER with either Venus-BirA* or BirA*-Bcl-XL were incubated in 50µM biotin for 18 hours, in the presence of either DMSO, 4-OHT or ABT-737. Biotin labelled proteins were isolated by streptavidin-affinity purification, then immunoblotted for vinculin, ER, myc-tag and biotin. Whole lysates of DMSO treated cells were included as controls. Right hand panel displays ER band intensity quantified by infrared imaging imaging using an Odyssey CLx. Data show three independent replicates. Significance calculated by ANOVA, followed by Tukey’s multiple comparison test. ns = non-significant,* = p<0.05, **** = p<0.0001

MCF10A cells stably expressing BadER in combination with either Venus-BirA* or BirA*-Bcl-XL were generated. MCF10A cells do not express endogenous oestrogen receptor (ESR1, ER) and are not sensitive to oestrogen or oestrogen antagonists ^27^. To validate the experimental system, Venus-BirA*/BadER and BirA*-Bcl-XL/BadER MCF10A cells were treated with either 4-OHT or ABT-737, and immunostained for either the myc-tagged BirA* or ER (**Fig.4B**, **Fig.S4A**). 4-OHT, but not ABT-737, induced BadER colocalisation with mtHSP70 in both BirA*-Bcl-XL and Venus-BirA* cells. To confirm that BadER and ABT-737 induced mitochondrial priming we quantified apoptosis using an annexin V live-cell imaging assay (**Fig.4C**). Neither ABT-737 nor 4-OHT treatment induced apoptosis in cells expressing BadER, alone or co-expressing Venus-BirA* or BirA*-Bcl-XL. Etoposide induced apoptosis in all the lines, but significantly less in cells expressing BirA*-Bcl-XL. In all three lines treatment with either ABT-737 or 4-OHT sensitised cells to etoposide, showing that BadER activation or ABT-737 increased mitochondrial priming. BirA*-Bcl-XL/BadER cells showed greater sensitisation to apoptosis with ABT-737 than with 4-OHT. Finally, to confirm that 4-OHT induced BadER binding to BirA*-Bcl-XL, we cultured Venus-BirA*/BadER and BirA*-Bcl-XL/BadER cells in 50 µM biotin, and treated them with either DMSO, 4-OHT or ABT-737. Biotinylated proteins were isolated by streptavidin affinity purification, immunoblotted for ER, myc and biotin, and quantified using an Odyssey CLx system (**Fig.4D**). Biotin conjugated BadER was precipitated from the BirA*-Bcl-XL cell lysates, showing they interacted. BadER biotin labelling in BirA*-Bcl-XL cells was increased by 4-OHT but abolished by ABT-737 (**Fig.4D**). BadER did not bind to Venus-BirA* under any of the conditions.

Three independent replicates of Venus-BirA*/BadER and BirA*-Bcl-XL/BadER MCF10A cells were treated with DMSO, 4-OHT or ABT-737, labelled with biotin and analysed by LC-MS/MS. Just under 2000 proteins were identified in each condition, with approximately 30 preys per bait reaching the BFDR threshold compared with Venus-BirA* (**Fig.5A**). Excluding Bcl-XL itself, 18 preys had a BFDR ≤0.05 in all three conditions (**Fig.5B, Fig.S5A**). We compared the average intensity among DMSO, 4-OHT and ABT-737 treated cells for all prey proteins with a BFDR ≤0.05 for at least one condition and subjected the results to hierarchical cluster analysis (**Fig.5C**). Many prey proteins showed distinct enrichment patterns between unprimed cells and those primed with 4-OHT or ABT-737. As expected, Bad and oestrogen receptor (ESR1) were both isolated from DMSO treated cells, were further enriched by 4-OHT and inhibited by ABT-737, confirming that BadER and ABT-737 competitively bind Bcl-XL. Of canonical interactions, endogenous Bid was enriched in unprimed (DMSO) cells, but this was reduced by BadER and ABT-737, suggesting that Bid is competed off by both FL Bad and the mimetic. Bcl-2 was identified under all three conditions, was enriched in cells primed by BadER, but reduced by ABT-737, suggesting that a Bcl-XL/Bcl-2 complex was stabilised by FL Bad binding, but disassembled by the Bad BH3-mimetic. Thus, canonical Bcl-XL interactions are detected, but enrichment is dependent upon whether cells are primed by a FL BH3-only protein or BH3-mimetic.

**Figure 5.**
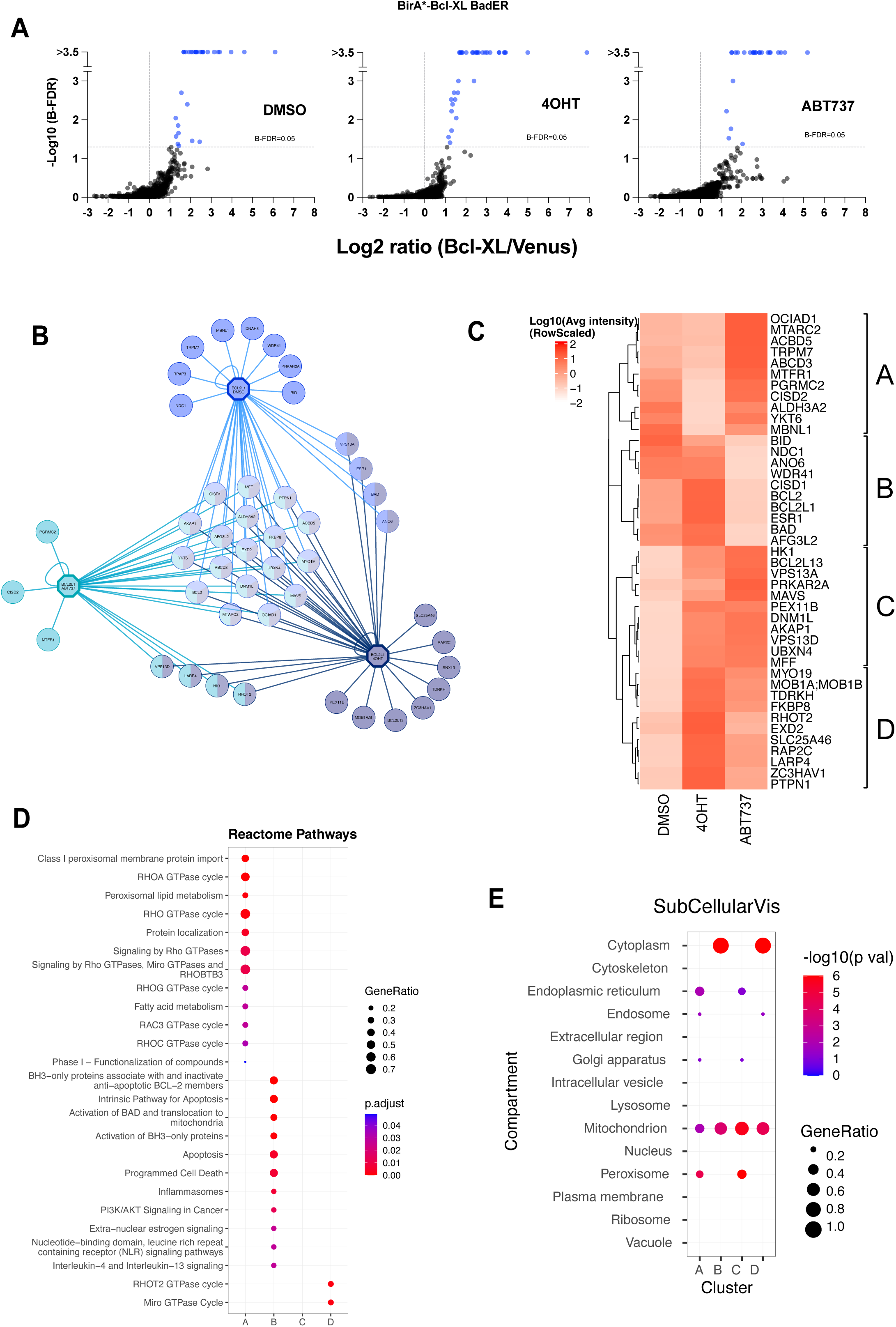
The BirA*-Bcl-XL proximity interactome shows differential changes when cells are primed by either Bad or the Bad BH3-mimetic, ABT-737. **A.** SAINT analysis identifies prey proteins labelled in cell expressing BirA*-Bcl-XL/BadER relative to those expressing Venus-BirA*/BadER, under treatment with DMSO, 4-OHT (to activate BadER), or ABT-737. Similar numbers of total identified proteins and significant preys (B-FDR <0,05) were identified in each condition. **B.** Significant BioID preys identified in XL-BirA*/BadER cells under each treatment condition (DMSO, 4-OHT and ABT-737) and their overlap. **C.** Heatmap displaying the relative enrichment of prey proteins across the DMSO, 4-OHT and ABT-737 conditions. Prey proteins where BFDR <0.05 in at least one treatment conditions are included. Hierarchical clustering revealed distinct enrichment patterns, enabling grouping into four clusters (A-D). A - enriched with ABT-737, decreased with 4-OHT; B - enriched with 4-OHT, decreased with ABT-737; C and D - enriched by either 4-OHT or ABT-737. Of note, preys identified that were in both clusters C and D did not show significant enrichment in unprimed (DMSO treated) cells. Data represents the average intensity from three independent replicates and have been scaled by row for visualisation. **D.** Over-representation analysis of prey clusters identified in Fig. 5C, based on Reactome annotations. **E.** Over-representation analysis of prey clusters identified in Fig. 5C, based on Cellular Component gene ontology terms.

Most prey proteins were non-canonical and showed distinct patterns of enrichment depending on whether cells were primed by 4-OHT-activated BadER or ABT-737 (**Fig.5C**). Previously described non-canonical interacting proteins that showed priming-dependent enrichment included Drp1 (DNML1) and FKBP8 ^11,12^. We also identified defined protein complexes, predicted through STRING (**Fig.S5B,C**), such as AKAP1 and its binding partner cAMP-dependent protein kinase type II regulatory subunit (PRKAR2A) ^28^. We divided the heat map into four prey groups: A, those inhibited by BadER; B, inhibited by ABT-737; C and D, poorly labelled in unprimed cells but differentially enriched by BadER and ABT-737. We compared these groups using gene ontology analysis (**Fig.5D,E**). Groups A and D were enriched for Reactome pathways associated with GTPase signalling, including those linked to intracellular membrane dynamics and peroxisomes. Group B was dominated by terms linked to cell death and apoptosis. Group C were not enriched for terms associated with Reactome pathways, but analysis of Cellular Component ontology terms indicated enrichment for peroxisomes, endoplasmic reticulum, Golgi and mitochondria (**Fig.5E**). Group C was particularly enriched for OMM proteins under primed condition, including hexokinase (HK1), MAVS, AKAP1, Drp1 and MFF, as well as proteins associated with mitochondrial contact sites and mitophagy.

We asked how the Bcl-2 interactome was affected by ABT-737 or BadER. MCF10A cells expressing BirA*-Bcl-2 and BadER were generated (**Fig.S6A**). We validated expression by immunofluorescence, immunoblotting, and the ability of ABT-737 or 4-OHT to reverse BirA*-Bcl2 dependent suppression of apoptosis (**Fig.S6B**). We then performed BioID analysis as before (**Fig.S6C**). As with BirA*-Bcl-XL, several proteins showed enrichment with BadER and/or ABT-737 priming. Some preys, such as AKAP1, YKT6 and PTPN1, were common with BirA*-Bcl-XL. However, Bcl-2 showed priming-dependent enrichment for nuclear envelope components, such as BANF1 and emerin, proteins within the ER (the chloride channel CLCC1), and CKAP4. Interestingly, prey proteins common between BirA*-Bcl-XL and BirA*-Bcl-2 showed similar enrichment patterns following priming. For example, where Bcl-2 was enriched when BirA*-Bcl-XL/BadER MCF10A cells were treated with 4-OHT, Bcl-XL (BCL2L1) was enriched in the same conditions in the BirA*-Bcl-2/BadER line. For both baits, AKAP1 was enriched with both 4-OHT and ABT-737, but YKT6 was enriched only by the mimetic.

These data suggest a complex set of interactions with anti-apoptotic Bcl-2 proteins on multiple subcellular membranes, with both canonical and non-canonical partners, which are differentially regulated when cells are primed by FL Bad or a Bad BH3-mimetic.

### Prey proteins indicate widespread priming-dependent changes with multiple intracellular membrane compartments and metabolism

Our BioID data suggested that interactions between anti-apoptotic proteins and novel partners were associated with changes in apoptotic priming. We previously reported that Bcl-XL retrotranslocation from mitochondria is reduced when bound to BH3-only proteins ^7^. We hypothesised that this reduced retrotranslocation might explain enrichment for membrane associated prey proteins by BioID. We therefore asked if BadER and ABT-737 both reduced Bcl-XL retrotranslocation. MCF10A cells expressing BadER and GFP-Bcl-XL were assessed by fluorescent recovery after photobleaching (FRAP) following treatment with either 4-OHT or ABT-737 (**Fig.6A,6B**). Although 4-OHT activation of FL BadER reduced GFP-Bcl-XL FRAP, consistent with our previous data, ABT-737 had no effect on retrotranslocation. As with the BioID results, these data indicate FL Bad and ABT-737 initiate distinct interactions with Bcl-XL on membranes, resulting in different downstream responses.

**Figure 6.**
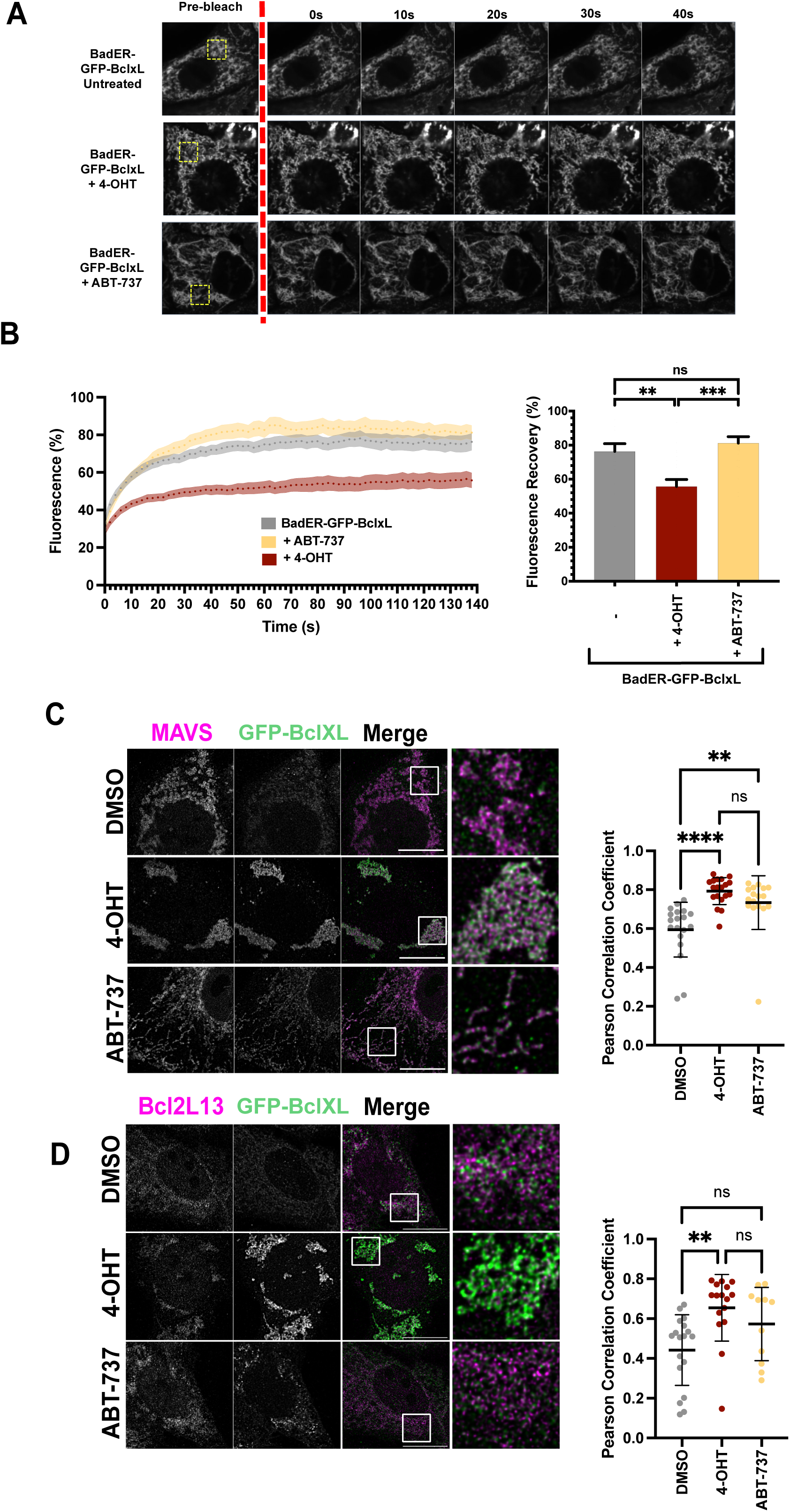
Full-length Bad and the Bad mimetic ABT-737 show differential effects on Bcl-XL membrane localisation. **A.** MCF10A cells expressing BadER and GFP-Bcl-XL were subjected to fluorescence recovery after photobleaching (FRAP) analysis as described previously ^7^ and in the methods. FRAP experiments were carried out with cells treated with either DMSO, 10 µM 4-OHT or 5 µM ABT-737. Cells were photobleached within the region of interest (ROI) indicated by the yellow box, and imaged post-bleach recovery for 140 seconds. **B.** Quantification of FRAP analysis from Fig6A. Fluorescence recovery was measured in the bleached ROI and the percentage recovery calculated. Percentage fluorescence recovery displayed as mean +/- SEM. Right hand panel: Fluorescence recovery at 140s post-bleach as a percentage of the original fluorescence. n = >20 cells per condition. Analysed by one-way ANOVA. **C.** STED imaging of MCF10A cells expressing BadER and GFP-Bcl-xL. Cells were immunostained with anti-MAVS and anti-GFP. Cells were treated with DMSO, 4-OHT or ABT—737 for 18 hours before fixing and immunostaining. Pearson correlation coefficient analysis was performed to measure co-localisation between MAVS and GFP, analysed by one-way ANOVA with multiple comparisons. **D.** STED imaging as in Fig6C, with cells immunostained for anti-GFP and anti-BCL2L13. Statistical analysis performed as described in Fig. 6C.

To validate some of the priming-dependent interactions with Bcl-XL, we carried out stimulated emission depletion (STED) microscopy. MCF10A cells expressing BadER and GFP-Bcl-XL were treated with either 4-OHT or ABT-737, fixed and immunostained for either endogenous MAVS or BCL2L13, along with anti-GFP. We chose these two preys as they were enriched by both 4-OHT and ABT-737, and we had good primary antibodies for STED imaging. In agreement with the FRAP data, more GFP-Bcl-XL was associated with mitochondria following BadER activation ^7^, but this was not apparent in cells treated with ABT-737. Despite this, Pearson correlation coefficient analysis confirmed that both MAVS **(Fig.6C)** and BCL2L13 **(Fig.6D)** co-localisation with GFP-Bcl-XL increased with either 4-OHT or ABT-737. We also noted that BadER activation resulted in increased clustering of mitochondria, seen with GFP-Bcl-XL.

Given the number of non-canonical proximity interactions identified for Bcl-XL and Bcl-2 we wondered whether distinct biological processes might be altered in cells under the different priming conditions. To assess this, whole protein lysates from BadER/BirA*-Bcl-XL MCF10A cells treated with DMSO, 4-OHT or ABT-737 were analysed by quantitative LC-MS/MS (**Fig.7A**). Whilst much of the proteome remained unchanged between conditions, hierarchical cluster analysis indicated key differences between 4-OHT and ABT-737-treated cells **(Fig.7B)**. Notably, there was little overlap between protein clusters both up and down regulated by ABT-737 or 4-OHT. GO analysis for biological processes indicated that clusters 2, 4 (downregulated by 4-OHT) and 7 (upregulated by 4-OHT) link FL-BadER with lipid, cholesterol and alcohol metabolism as well as downregulated mitochondrial gene expression and translation **(Fig.7C)**. In contrast, ABT-737 downregulated pathways associated with RNA metabolism and nuclear organisation (cluster 5) and upregulated the endoplasmic-reticulum-associated protein degradation (ERAD) pathway, protein folding, endoplasmic reticulum stress and glycosylation (cluster 6). Individual proteins from these pathways were showed differential expression. In ABT-737 treated cells, upregulated proteins included PDIA4, HYOU1, HSPA5 (binding immunoglobulin protein, BiP) and HSP90B1, all involved in the endoplasmic reticulum stress response. In 4-OHT-treated cells, proteins linked to lipid metabolism (LDLR, DHCR24), lysosome function (PPT1, ASAH1) and mitophagy (SQSTM1/p62) were altered.

**Figure 7.**
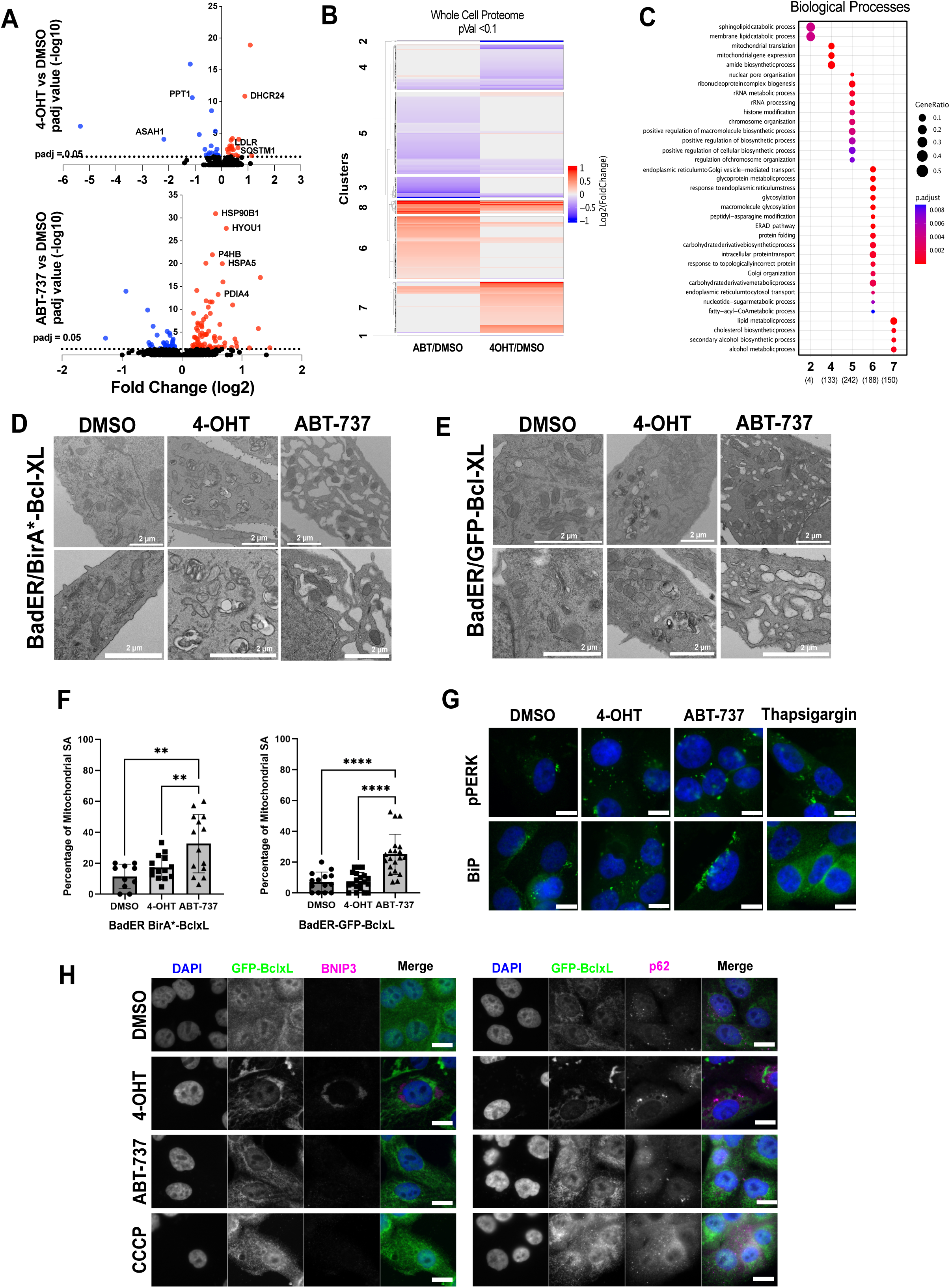
Full-length Bad and the Bad mimetic ABT-737 have distinct effects on mitochondrial and ER function. **A.** Whole-cell proteomic analysis of MCF10A cells expressing BadER and BirA*-Bcl-XL following 18 hour DMSO, 4-OHT or ABT-737 treatment. Fold change represent 4-OHT or ABT-737 treated cells relative to DMSO treated cells. Data represent the mean values from three independent replicates. P-values were adjusted for multiple comparisons using Benjamini-Hochberg (BH) correction. **B.** Heatmap displaying data from Fig.7A, showing differential abundance of proteins in 4-OHT or ABT-737 treated cells relative to DMSO control. Only proteins with P-value >0.1 are displayed. Hierarchical clustering identifies eight clusters (1-8). **C.** Over-representation analysis of protein clusters identified in Fig. 7B., based on Biological Processes gene ontology terms. Terms over-represented by proteins in clusters 2, 4, 5, 6 and 7 show distinct cellular processes affected by BadER activation and ABT-737. **D.** Transmission electron microscopy of BadER-GFP BirA*-Bcl-xL MCF10A cells following 18 hours treatment with DMSO, 4-OHT or ABT-737. In 4-OHT treated cells, clusters of mitochondria are indicated by the red arrows, and mitophagic vesicles by the yellow arrows. Examples of mitochondrial endoplasmic reticulum contacts (MERCs) are indicated by the green arrows. Scale bar = 2µm. **E.** Transmission electron microscopy of BadER-GFP-Bcl-xL MCF10A following 18 hours treatment with DMSO, 4-OHT or ABT-737. In 4-OHT treated cells, clusters of mitochondria are indicated by the red arrows, and mitophagic vesicles by the yellow arrows. MERCs are indicated by the green arrows. Scale bar = 2 µm. **F.** Mitochondria-endoplasmic reticulum contact site analysis for both cell lines. Contact sites were measured as a percentage of mitochondrial surface area across multiple TEM images. Each data point = one image. Analysis was carried out by one-way ANOVA with multiple comparisons. Significance = p<0.05. **G.** Immunofluorescence staining of BadER-GFP BirA*-Bcl-xL MCF10A for pPERK and BiP following treatment with DMSO, 4-OHT, ABT-737 or thapsigargin, as described in the methods. Scale bar = 10 µm. **H.** BadER-GFP-Bcl-xL MCF10A cells immunostained stained for anti-GFP and either BNIP3 or p62 following treatment with DMSO, 4-OHT, ABT-737 or CCCP, as indicated. Scale bar = 10 µm.

These data indicated profound differences in how FL Bad and ABT-737 might affect organelle organisation in non-apoptotic cells. To investigate this further, we examined BadER/BirA*-Bcl-XL cells treated with either DMSO, 4-OHT or ABT-737 by electron microscopy (EM) **(Fig. 7D)**. Cells primed by full-length BadER showed clusters of healthy mitochondria as well as what appeared to be mitophagic clearance of others. Mitophagy was not apparent in cells treated with ABT-737, but instead there were striking changes in the endoplasmic reticulum network, which appeared swollen with increased mitochondrial contacts. To confirm this, we undertook EM imaging of BadER/GFP-Bcl-XL cells **(Fig.7E)**. These also showed increased mitophagy following BadER activation, with pronounced clustering of healthy mitochondria. Again, ABT-737 resulted in endoplasmic reticulum swelling. Quantification of mitochondrial-endoplasmic reticulum contact surface area confirmed that ABT-737 significantly increased the proportion of contact sites compared to both DMSO and 4-OHT-treated cells **(Fig.7F).** The increase in mitochondria-endoplasmic reticulum contacts is consistent with enrichment of several membrane contact site prey proteins in the BioID analysis.

Together, the BioID, whole cell proteomics and EM analysis indicated that ABT-737 and FL BadER had broad but distinct effects beyond canonical mitochondrial priming. To confirm the changes seen in the EM were indicative of ER stress, we performed immunofluorescence analysis for markers of the unfolded protein response (UPR) **(Fig.7G)**. There was an increase in both phosphorylated PERK and BiP (HSPA5), one of the most significantly upregulated proteins identified in the proteomics analysis **(Fig.7A)**, in cells treated with either ABT-737 or thapsigargin (a positive control to induce ER stress), but not with 4-OHT, compared with DMSO treated cells. We also examined the mitophagy markers BNIP3 and p62 (also identified as significant in the proteomics analysis; SQSTM1) by immunofluorescence **(Fig.7H)**. Consistent with the EM data, cells primed with BadER showed increased BNIP3 and p62 foci, although the latter was more pronounced in cells treated with the mitochondrial uncoupler carbonyl cyanide m-chlorophenyl hydrazone (CCCP, positive control for mitophagy). Thus, mitophagic signalling may be initiated by full-length Bad, whereas ABT-737 appears to induce endoplasmic reticulum stress.

## Discussion

Here we describe a BioID strategy to interrogate the wider Bcl-2 protein family interactome in non-apoptotic cells. By utilising multiple Bcl-2 protein baits, combined with inducible mitochondrial priming, we defined a dynamic proximity interactome regulating diverse downstream processes. Although components of the canonical Bcl-2 interactome were detected, most preys identified by Bcl-XL, Bcl-2 and Bax lay beyond their defined literature curated networks. Some preys, such as Drp1 and FKBP8, are interacting partners of Bcl-2 proteins in the literature ^11,12^. Other preys included known multiprotein complexes, including VPS13D and RHOT2, which link ER, mitochondria and peroxisomes ^29^, and the mitochondrial adaptor AKAP1 and cAMP-dependent protein kinase regulatory subunit PRKAR2A ^28^. These proximity partners defined membrane contacts as foci for priming-dependent changes and suggest wider regulation of cellular processes by Bcl-2 proteins. Indeed, although FL Bad and the Bad mimetic ABT-737 both increased mitochondrial priming, they enriched for distinct subsets of preys for Bcl-2 and Bcl-XL as well as driving different downstream effects. Thus, whereas BadER brought about changes focussed at mitochondria, the BH3-mimetic altered protein transport and induced an endoplasmic reticulum stress response.

Current models of Bcl-2 family regulation focus on canonical heterodimeric interactions defined largely around commitment to MOMP ^1^. Permutations of these interactions define distinct modes of apoptosis regulation ^30^. However, most cells exist in an unprimed state and interactions of Bcl-2 proteins in this context remain ill defined. Our BioID data identifies canonical interactions between the three subgroups of Bcl-2 protein but highlight distinctions from the accepted models of their regulation. Bid-BirA* identified four significant preys, of which three were known binding partners, Bcl-XL, Bcl-2 and Bax. Interestingly, Bid-BirA* is full-length and we have no evidence of significant caspase cleavage of it in our cells. Therefore, caspase cleavage is not essential for Bid binding to canonical partners. BirA*-Bcl-XL labelled four Bcl-2 family preys: itself, although not possible to distinguish between self-labelling and endogenous Bcl-XL; Bcl-2; Bad, but again not possible to distinguish between BadER and endogenous Bad; and endogenous Bid. These data indicate that in non-apoptotic cells, in the absence of directly activating caspase 8, full-length Bid is the only endogenous BH3-only protein unambiguously identified as a canonical Bcl-XL binding partner. Previous studies have shown that Bid is functional in the absence of caspase 8 cleavage ^31,32^, and it was originally identified through the ability of its full-length form to bind directly to Bcl-2 and Bax ^33^. Taken together, the data suggest that in the absence of caspase cleavage Bid functions as a sensitiser BH3-only protein, and not exclusively as a direct activator.

Of other canonical interactions, BirA*-Bcl-XL and BirA*-Bcl-2 labelled endogenous Bcl-2 and Bcl-XL respectively, suggesting multiprotein complexes of these at membranes and more complex regulation than in generally accepted models. We have previously shown that mitochondrial Bcl-XL is almost exclusively in a high molecular weight complex in non-apoptotic cells ^34^, and Bogner et al. described higher order complexes of Bcl-XL that inhibited proapoptotic Bcl-2 proteins whilst simultaneously binding multiple BH3-proteins ^35^. These Bcl-XL complexes concurrently bound both Bid and Bad and facilitated allosteric activation between them, distinct from the accepted model of BH3-protein displacement. A more complex model of Bcl-2 interactions is also supported by our BioID data in the presence of conditional activation of BadER or the Bad BH3-mimetic, ABT-737. When BadER was activated, both BirA*-Bcl2 and BirA*-BclXL labelled more endogenous Bcl-XL and Bcl-2 respectively, indicating it promoted complex formation. BadER also reduced labelling of endogenous Bid by BirA*-Bcl-XL. In contrast, ABT-737 almost completely abolished Bid binding to BirA*-BclXL, but also significantly reduced the labelling of endogenous Bcl-2 and Bcl-XL. These data suggest very different effects of full-length Bad and the Bad mimetic on canonical Bcl-2 protein complexes, which was supported by the FRAP measurements of GFP-Bcl-XL dynamics. Thus, although both ABT-737 and BadER both primed cells for apoptosis and displaced endogenous Bid from Bcl-XL, BadER stabilised multiprotein complexes of Bcl-2 and Bcl-XL whereas ABT-737 led to their disassembly. These data align with other studies showing BH3 mimetics differ from full-length proteins ^25,26^. Full-length BH3-only proteins have interactions independent of their BH3-domains which may promote the assembly of multiprotein complexes at membranes that are not seen with mimetics. This raises questions about the role of these complexes in priming and other functions at mitochondria.

We found multiple, priming-dependent interactions that fall outside the canonical Bcl-2 family network and which suggested functions beyond apoptosis. Despite numerous reports of non-canonical interactions with Bcl-2 proteins, a holistic approach to this wider interactome has been missing until now. Our data identify previously described interacting partners, such as FKBP8 which recruits Bcl-2 proteins to mitochondria ^12^. Elements of the mitochondrial fission machinery, with reported roles in Bcl-2 protein function, were also identified. Drp1 (DNML1) was a robust prey, identified by Bcl-XL, BaxS184V and Bcl-2, and enriched in primed cells. MFF, another component of the mitochondrial fission machinery, was identified by BirA*-Bcl-XL. Hexokinase 1 (HK1) was another prey in the same BirA*-Bcl-XL cluster as Drp1 and MFF. During the preparation of this manuscript, HK1 was shown to compete with Drp1 for MFF binding under conditions of nutrient stress, inhibiting mitochondrial fission ^36^. Thus, our data can be predictive of new components of protein complexes, identifying novel regulatory pathways that interface with the apoptotic machinery. Other preys also included known components of multi-protein complexes, including AKAP1 and PRKAR2A. AKAP1 was another robust prey enriched in primed cells, identified by Bcl-2, Bax and Bcl-XL baits. Another robustly identified prey was MAVS, which regulates cellular responses to viral dsRNA ^37^. MAVS activation results in its assembly into an oligomerised signalling platform to initiate the innate immune response through type-1 interferons, NF-κB and apoptosis. We did not see evidence that priming alone induced interferon expression (data not shown), but did see increased co-localisation of MAVS with GFP-Bcl-XL by STED. MAVS mitochondrial puncta form in response to dsRNA and are associated with apoptotic cell death ^38^. One possibility is that MAVS interfaces with the apoptotic machinery but requires dsRNA to fully initiate downstream effects.

Analysis of enriched GO terms implicated diverse biological processes that may regulate or be regulated by Bcl-2 proteins, and which were distinct between priming by ABT-737 or BadER. BadER resulted in changes more focused at mitochondria. In contrast, ABT-737 with BirA*-Bcl-XL enriched for endoplasmic reticulum and peroxisome components, whilst BirA*-Bcl-2 labelled components on multiple membranes, including the endoplasmic reticulum and nuclear envelope. Given the known promiscuity of Bcl-2 ^39^, these hits verify its localisation to a range of organelles and roles in processes such as calcium homeostasis ^40,41^ and transcription factor trafficking ^42,43^. Analysis of the whole cell proteome indicated reorganisation of mitochondria in response to BadER, whereas ABT-737 impacted on protein transport, Golgi organisation and the UPR. EM confirmed this, with BadER-dependent priming initiating mitophagy, supported by BioID enrichment of BCL2L13, where ABT-737 resulted in a significant reorganisation of the endoplasmic reticulum, with a notable increase in mitochondrial contact sites. It will be important to identify the key interaction partners mediating these diverse responses, although there are interesting candidates, such as the anion channel CLCC1, a prey identified by BirA*-Bcl-2. CLCC1 variants are linked to amyotrophic lateral sclerosis, and its knockout leads to endoplasmic reticulum stress, misfolded protein accumulation and a swollen endoplasmic reticulum phenotype like that seen in cells treated with ABT-737 ^44^.

The number of BioID preys identified associated with membrane contact sites was striking. These included VPS13D and RHOT2 (also known as Miro), which tether mitochondria, endoplasmic reticulum and peroxisomes ^29^. The VPS13D-RHOT2 complex was only significantly detected when cells were primed with either ABT-737 or BadER. VPS13D has multiple roles at contacts sites, including lipid transfer and ubiquitin-mediated mitochondrial clearance ^45^. Other known contact site proteins identified included VPS13A ^29^, PRKAR2A ^22^, EXD2 ^46^, FKBP8 ^22,46,47^, MYO19 ^48^, PTPN1 and the AKAP1-DNM1L-MFF complex ^22^. Interestingly, a recent study mapped the mitochondrial proximity interactome by BioID, using 100 mitochondrial baits across all compartments (but no Bcl-2 family members) ^22^. Several of the OMM baits used in that study were identified in our current analysis as significant prey proteins, including AKAP1, Drp1, FKBP8, MAVS and OCIAD1. As baits, these proteins identified several membrane contact site components shared between their data set and ours, but they did not identify any Bcl-2 family preys. However, in our results these proteins were all enriched in conditions where cells were primed for apoptosis. Antonicka et al. used HEK293T cells in complete medium, making it is unlikely that those cells could be considered “primed”, so we might not expect Bcl-2 protein interactions to be enriched. They did identify the same regulatory subunits of the PKA complex, indicating that these sites may coordinate kinase signalling and priming. Together, these data indicate that membrane contact site composition may be modified in response to changes in apoptotic priming, highlighting the potential importance of these sites as mitochondrial signalling hubs. It will be important to understand how priming alters contact site functions, such as lipid exchange or organelle dynamics.

Overall, there are now significant data to indicate that Bcl-2 proteins function within a complex milieu of OMM proteins, membrane contacts and dynamics. This suggests a potential for Bcl-2 proteins to influence and be influenced by a range of cellular signals and physiological states. Understanding this wider influence will provide important insights into how apoptosis is coordinated.

## Limitations of this study

BioID and mass spectrometry approaches have well-known limitations, in particular the ability to label proximity partners based upon availability of lysine residues and not detecting tryptic peptides for some preys. This means that some expected preys may not be detected. Bax-S184V did not identify anti-apoptotic Bcl-2 proteins, for example. Some well characterised interactions, such as VDAC isoforms, were also not detected with our baits ^49^. Missing expected preys may also be due to specific cell type and experimental conditions. We previously identified VDAC2 using Bid-BirA*, but very specifically in cancer cells arrested in mitosis ^15^. The MCF10A cells used here were largely confluent, appropriate for epithelia, but with a negligible mitotic index. We might have expected to see VDAC2 with BirA*-Bax, although Bak might be a more suitable bait for examining this interaction ^50^. Indeed, further expansion of the Bcl-2 interactome with other family members will be interesting. A Bad BioID interactome was published recently, also using MCF10A cells ^51^. As they used cells in complete media, with a full complement of serum and growth factors, priming dependent interactions at the OMM might not be expected as Bad would be largely in a phosphorylated state. As expected, they identified several 14.3.3 isoforms, as well as numerous non-canonical partners.

Canonical Bcl-2 protein binding partners might be more associated in cells undergoing MOMP and apoptosis. We did not look at apoptotic cells in the present study as that brings problems for BioID. The process of apoptosis itself is relatively quick in an individual cell, occurring over a few minutes, beyond the scope of BioID labelling. Furthermore, as cells within a population die asynchronously, any labelling will be a biochemical average predominantly of cells that are yet to die, with a minor component from those dying. It is unlikely that any significant labelling would occur in cells post MOMP. Thus, meaningful analysis of and data from “apoptotic” cells would be difficult.

Finally, any approach using exogenously expressed proteins invites criticism around expression levels. One necessity for using exogenous expression here was to balance the levels of the baits and controls using tagBFP, essential for comparative quantitative analysis of the MS data. We mitigated exogenous expression in part with the comparison of different priming states within the same cell lines and baits. Developing approaches with endogenous BioID knock in baits will be an important future approach, but low and variable expression levels of some Bcl-2 proteins will present challenges around specificity and sensitivity of detection.

## Methods

### Cell culture and treatments

All MCF10A cell lines were cultured in DMEM/F-12 50/50 1x medium supplemented with 5% sterile filtered horse serum, 1% penicillin/streptomycin, 20ng/mL EGF, 0.5µg/mL hydrocortisone, 100ng/ml cholera toxin and 10 µg/ml insulin. All cells were incubated at 37°C with 5% CO_2_.

### Lentiviral generation of stable cell lines

To produce the lentivirus required to generate stably transduced cell lines, HEK293T cells were seeded to approximately 70% confluency, before being transfected with 3 µg MD2.G, 4.5 µg psPax2 and 6 µg of the relevant expression plasmid. This was mixed with PEI at a concentration of 1 µg/µL before being added to cells and incubated overnight. Transfection media was removed and replaced with 10 mL growth media supplemented with 100 μL (1% v/v) sodium butyrate (Sigma-Aldrich #19137), then incubated for 6–8 hours. This was then replaced with fresh growth media and left for 48 hours to allow cells to produce lentivirus. Lentivirus media was injected through a 0.45 µm filter and virus precipitated in 1x lentivirus precipitation solution at 4°C overnight. Precipitated lentivirus was pelleted by centrifugation for 30 mins at 1500xg, then resuspended in approximately 250 µL supernatant. Lentivirus was stored at −80°C.

To generate stable cell lines, MCF10A cells were seeded at an appropriate density in 6-well plates and left to adhere overnight. Media containing 8 µg/mL polybrene was added, and a lentivirus aliquot added dropwise. This was left for 48 hours before being replaced with growth media. Once confluent, cells were cultured to higher densities for approximately 14 days to allow dissipation of transient construct expression. Stably expressing cells were then trypsinised and pelleted before being resuspended in FACS media (serum-free DMEM-F12 supplemented with 1% penicillin/streptomycin and 25 mM HEPES). Cells were passed through a 50 µm filter and sorted using a FACS Aria Fusion (BD Biosciences).

### Western blotting

Western blotting was carried out according to previous protocols. Blots were visualised using a LI-COR Odyssey CLx imaging system. Images were processed and analysed using Image Studio Lite (v5.2). The following antibodies and bacterial proteins were used for Western blotting:

| **Antibody** | **Host** | **Concentration** | **Supplier** | **Catalogue Number** |
| --- | --- | --- | --- | --- |
| Anti-Vinculin | Mouse | 1:1000 | Sigma-Aldrich | V9131 |
| Anti-Myc-tag<br>Clone 4A6 | Mouse | 1:1000 | Millipore | 05-725 |
| IRDye® 680RD<br>Streptavidin |  | 1:10,000 | LI-COR | 926-68079 |
| Anti-ERα | Rabbit | 1:1000 | Abcam | ab16660 |
| IRDye® 800CW<br>anti-Mouse IgG | Donkey | 1:10,000 | LI-COR | 925-32212 |
| IRDye® 680RD<br>anti-Rabbit IgG | Donkey | 1:10,000 | LI-COR | 925-68073 |

### Biotin labelling and affinity purification of biotinylated proteins

Proximity labelling was induced and cells lysed as previously described ^15^ with the following modifications: cells were incubated in 50 µM biotin (Sigma-Aldrich) supplemented media for 18 hours; cells were lysed in RIPA supplemented with 1x protease inhibitor cocktail (Millipore).

Biotinylated proteins were isolated using streptavidin-agarose beads (Sigma-Aldrich). 50 µL of bead slurry was washed twice with lysis buffer and then pelleted by centrifugation. Volume equivalent to 500 µg total protein of cell lysate was added to beads and the volume made up to 500 µL with lysis buffer. Lysates and bead slurry were incubated overnight on an end-over-end tumbler at 4°C. The next day beads were pelleted, washed twice in lysis buffer, once in urea wash buffer (2 M urea, 10 mM Tris-HCl pH 8.0), and then three times with lysis buffer. Following the final wash, beads were resuspended in 50 µL 1x lithium dodecyl sulphate sample buffer (Novex, NuPAGE) supplemented with 2 mM biotin and 50 mM dithiothreitol (DTT) and incubated at 95°C for 5 minutes to elute bound proteins. 45 µL of eluted sample was immediately subjected to SDS PAGE and in-gel tryptic digestion in preparation for mass spectrometry, while the remaining 5 µL was analysed *via* western blotting to confirm biotinylation and successful protein enrichment.

### Sample preparation for liquid chromatography mass spectrometry

For BioID experiments, streptavidin affinity purified samples were briefly subjected to SDS-PAGE to run samples into the resolving gel, then fixed and stained using InstantBlue (Expedeon). Gel fragments containing protein samples were digested with trypsin and peptides extracted as described ^15^. For whole-cell lysate proteomics experiments, volumes of lysate equivalent to 50 µg total protein were prepared for LC-MS/MS using S-Trap columns (ProtiFi) as per the manufacturer’s protocols. Tryptic peptides were desalted using POROS Oligo R3 beads (Thermo-Fisher Scientific). Beads were prepared with 50% acetonitrile (v/v) in water, then peptides added. Beads/peptides were washed using 0.1% trifluoracetic acid (v/v) in water and then eluted with 50% acetonitrile (v/v) in water. Eluted peptides were dried via vacuum centrifugation and then resuspended in 5% acetonitrile (v/v), 0.1% trifluroacetic acid (v/v), in water.

### Mass spectrometry data acquisition

Tryptic peptides were subjected to liquid-chromatography tandem mass spectrometry (LC-MS/MS) using a 3000 Rapid Separation LC (Dionex Corp.) coupled to a Q Exactive HF mass spectrometer (Thermo Fisher Scientific). Mobile phase A was 0.1% (v/v) FA in water, mobile phase B was 0.1% (v/v) FA in ACN, and a 75 mm × 250 µm internal diameter 1.7 mM CSH C18 analytical column (Waters) was used. 3 µl of sample was transferred to a 5-µl loop and loaded onto the column at a flow rate of 300 nl/min for 13 min at 5% (v/v) mobile phase B. The loop was taken out of line and the flow was reduced to 200 nl/min in 30 s. Peptides were separated using a gradient of 5% to 18% B in 34.5 min, from 18% to 27% B in 8 min, and from 27% to 60% B in 1 min. The column was washed in 60% B for 3 min before re-equilibration to 5% B in 1 min. Flow was increased at 55 min to 300 nl/min until the end of the run at 60 min. Peptides were selected for fragmentation automatically by data-dependent analysis.

Raw data were processed using *MaxQuant* v1.6.2.3 against the human and/or mouse proteome obtained from Uniprot (July 2023) ^52^. Default parameters were used for analysis, in addition to the inclusion of lysine biotinylation as a variable modification, matching between runs and only using unique peptides for protein.

### Bioinformatics data analyses

#### Whole-cell lysate proteomics data analysis

Differential abundance analysis was performed using MSqRob ^53^.Treatment was considered a fixed effect, whereas peptide sequence was considered to be random effects.

#### BioID proteomics data analysis

*MaxQuant* derived Protein LFQ intensities were analysed by SAINTexpress ^54^ using default parameters to determine the confidence of bait-prey interactions. Preys with a BDFR value of ≤0.05 were considered to be high-confidence bait-prey interactions.

Heatmaps were generated through the ComplexHeatmap R package, with hierarchical clustering performed using Euclidian distances. Over-representation analysis of gene ontology terms was performed and visualised using the SubcellulaRVis (cellular component annotations) or clusterProfiler (all other annotations) packages in R ^55^. Interaction networks were visualised using Cytoscape [v3.9.1] ^56^. Predicted interaction networks were computed by STRING (Search Tool for the Retrieval of Interacting Genes/Proteins) ^57^. Only data from experimentally validated interactions and interactions in curated databases were used to score predicted functional partners. STRING database v11.0, accessed April 2020.

#### Immunofluorescence imaging

Samples were prepared according to previous protocols. Briefly, cells were fixed onto coverslips for 10 min in 4% v/v paraformaldehyde, then stained for targets (2-hour primary antibody incubation, 1-hour secondary antibody incubation). DAPI (Sigma-Aldrich #D9542) was added at a concentration of 1 μg/mL for 10 min before coverslips were washed, left to dry and mounted onto slides using Dako fluorescence mounting medium (Agilent #S3023). Immunofluorescence images were acquired using a Zeiss Axio Imager M2 fluorescence microscope. Images were captured using a 63x oil-immersion objective and a Hamamatsu ORCA-ER fluorescence digital camera with Micro-Manager software. Images were subsequently processed using *ImageJ software*. The following antibodies were used:

| **Antibody** | **Host** | **Concentration** | **Supplier** | **Catalogue Number** |
| --- | --- | --- | --- | --- |
| Anti-mtHSP70 | Mouse | 1:400 | Invitrogen | MA3-028 |
| Anti-Myc-tag<br>Clone 4A6 | Mouse | 1:400 | Millipore | 05-725 |
| Anti-ERα | Rabbit | 1:400 | Abcam | ab16660 |
| Anti-mCherry<br>[IC51] | Mouse | 1:400 | Abcam | ab125096 |
| Anti-Myc-Tag | Rabbit | 1:400 | Cell Signaling<br>Technology | 2272 |
| Anti-GFP | Rabbit | 1:400 | ChromoTek | pabg1 |
| Anti-pPERK<br>(Ser713) | Rabbit | 1:100 | BioLegend | 649401 |
| Anti-BiP | Rabbit | 1:100 | Cell Signaling<br>Technology | 3177 |
| Anti-BNIP3 | Mouse | 1:400 | Proteintech | 68091-1-Ig |
| Anti-p62 | Rabbit | 1:400 | Cell Signaling<br>Technology | 5114 |
| Anti-Mouse IgG<br>(H+L), Alexa<br>Fluor™ 555 | Goat | 1:400 | Invitrogen | A28180 |
| Anti-Rabbit IgG<br>(H+L), Alexa<br>Fluor™ 647 | Donkey | 1:400 | Invitrogen | A31573 |

#### Live-cell imaging apoptosis assays

Cells were treated with 800 µM etoposide (Sigma-Aldrich #E1383-25MG). 10 µM 4-OHT (Sigma-Aldrich #H7904) and 5 µM ABT-737 (APExBIO #A8193) were used. All drugs and vehicles were dissolved in phenol-red free complete MCF10A media supplemented with 2.5 mM CaCl_2_●2H_2_O, 2µL/mL 500x FITC Annexin V (Abcam #14082) and 2 µL/mL PI (Abcam #14083). Imaging was set up on an EVOS M7000 microscope. Cells were imaged every hour for 18 hours whilst being incubated at 37°C with 5% CO_2_ and humidity. A 10x objective with 0.3 aperture was used. Images were processed using *ImageJ*. A representative field of view was selected and tracked through time to cumulatively count the number of cells undergoing apoptosis.

#### Fluorescence Recovery After Photobleaching (FRAP)

FRAP was performed as previously described ^7^. Cells were washed with PBS before live-imaging medium (phenol red-free complete media supplemented with 25 mM HEPES) containing treatment (10 µM 4-OHT or 5 µM ABT-737) was added. Dishes were incubated in the microscope chamber at 37°C. Images were captured using a 3i Zeiss confocal system with CSU-X1 spinning disc (Yokogawa). A 63x/1.40 Plan Apochromat objective (Zeiss), an Evolve EM-CDD camera (Photometrics), and a motorised XYZ stage (Intelligent Imaging Innovations) driven by Marianas hardware were used. Slidebook 6.0 software was used to capture images.

### STED Microscopy

Cells were seeded onto coverslips for the appropriate drug treatments. After treatment, samples were rinsed three times in PBS, then fixed in 2% PFA for 15 min, followed by three more rinses in PBS. Samples were permeabilised in 0.1% Triton for 10 min, rinsed in PBS three times for 5 min each, then coverslips were blocked for 1 hour in 2% BSA. Coverslips were incubated for 1 hour with primary antibodies diluted in PBS/2% BSA, followed by three PBS washes of 5 min each. Coverslips were incubated for 1 hour with secondary antibodies diluted in PBS/2% BSA, washed in PBS three times, rinsed in ddH_2_O and left in the dark to completely dry before being mounted onto slides using ProLong™ Gold Antifade Mountant (Invitrogen™ #P36930).

STED images were acquired using a Leica TCS SP8 AOBS inverted gSTED microscope using a 100x/1.4NA HC PL APO (Oil; STED WHITE) objective and 2x confocal zoom. The confocal settings were: pinhole 1 airy unit, scan speed 400Hz unidirectional, format 2048 x 2048. STED images were collected using hybrid detectors with the following detection mirror settings: GFP 498–582nm; far red 557–765nm. STED images were collected sequentially. Leica LAS X v1.8.0.13370 software was used. Acquired images were analysed and the Pearson correlation coefficient calculated between the relevant channels of the multi-channel images to calculate the overlap between GFP-Bcl-xL and the target proteins.

The following antibodies were used:

| **Antibody** | **Host** | **Concentration** | **Supplier** | **Catalogue Number** |
| --- | --- | --- | --- | --- |
| Anti-MAVS | Rabbit | 1:200 | Proteintech | 14341-1-AP |
| Anti-BCL2L13 | Rabbit | 1:200 | Proteintech | 16612-1-AP |
| Anti-GFP Clone 1E10H7 | Mouse | 1:200 | Proteintech | 66002-1-Ig |
| Anti-Mouse IgG (H+L), Alexa Fluor™ 488 | Goat | 1:200 | Invitrogen | A-11001 |
| Anti-Rabbit IgG (H+L), Alexa Fluor™ 647 | Donkey | 1:200 | Invitrogen | A31573 |

### Electron microscopy

Cells were fixed with 2.5% glutaraldehyde and 4% formaldehyde in 0.1 M HEPES (pH 7.2) overnight. Fixed cells were scraped from petri dishes and centrifuged to form pellets with Eppendorf microcentrifuge at 7,000 rpm for 10 min. Cell pellets were post-fixed in reduced osmium (1.5% potassium ferrocyaninde, 1% osmium tetroxide in 0.1 M cacodylate buffer, pH 7.2) for 1 hour and in 1% uranyl acetate in water overnight. Samples were dehydrated in an ascending series of ethanol and embedded into TAAB Low Viscosity resin. Ultrathin sections of 70 nm were cut using a Leica UC7 ultramicrotome and mounted on 400 mesh copper grids. Images were acquired using a Thermo Fisher Talos L120C microscope at 120 kV high tension with a Ceta CMOS camera.

MERC contact site analysis was carried out in Fiji using a surface intersection method. Briefly, this involved overlaying a 1 µm calibrated square grid over each image and counting the number of membrane intersections with lines of the grid. The number of MERC intersections with lines of the grid was measured and presented in a graph as a percentage of the total mitochondria outer membrane grid line intersections.

### Live apoptosis assay

Cells were seeded at appropriate densities at least 24hrs before priming with DMSO, 10 μM 4-OHT or 5 μM ABT-737. After 18 hours, media was replaced by phenol-red free complete MCF10A media supplemented with 2.5 mM CaCl_2_●2H_2_O, 2 µL/mL 500X FITC Annexin V, 2 µL/mL PI. Imaging was carried out on an Incucyte system, where cells were incubated at 37°C, 5% CO_2_ and humidity. Images were taken at 10x magnification every hour for 23 hours. Images were analysed using ImageJ and the cumulative number of cells undergoing apoptosis were counted within the same fields of view over the time course.

## Acknowledgements.

RP was generously supported by a studentship funded by John and Janet Hartley, and post-doctoral positions funded through the Wellcome Trust Institutional Strategic Support Fund and CRUK Alliance for Cancer Early Detection (ACED). CELM, MJ, and LK were funded through the Manchester Cancer Research Centre by CRUK training awards. IT-H was supported by the Wellcome Trust PhD Programme in Quantitative and Biophysical Biology. The Wellcome Centre for Cell-Matrix Research is supported by a core grant from the Wellcome Trust (203128/Z/16/Z). We are grateful for the assistance provided by staff in the Bioimaging, Flow Cytometry and BioMS core facilities at the University of Manchester. The Bioimaging Facility microscopes used in this study were purchased with grants from BBSRC, Wellcome Trust and the University of Manchester Strategic Fund.

## Author contributions

Conceptualisation, RP, CELM, LEK, APG; Methodology and experimental work, RP, CELM, ELK, MJ; Data analysis, RP, CL, IT-H, AM; Data curation, RP, APG; Writing, RP, CELM, APG; Funding and project management, APG.

