## Supplemental data figures for "Multiplexed proximity labelling proteomics identifies a non-canonical Bcl-2 family interactome associated with apoptotic priming"

**Supplementary data figures**

**Figure S1. BioID identification of Bcl-2 protein family interactions in non-apoptotic cells.**

**A.** Representative images of immunostained DKO MEFs expressing mCherry-tBid, either alone or in combination with Venus-Bax or BirA*-Bax as described in 1B. Scale bar represents 10 mm.

**B.** Representative images of immunostained WT MEFs expressing mCherry-tBid either alone, or in combination with either Venus-Bcl-XL or BirA*-Bcl-XL as described in 1B. Scale bar represents 10 mm.

**C.** Venn diagram of gene lists derived from conditions described in 1E indicating limited overlap between predicted and observed interactions.

**D**. BioID observed Bid, Bax, Bcl-XL network extended with STRING-predicted interacting genes. The BioID observed Bid, Bax, Bcl-XL network (**Figure 1E**) was extended to display the STRING predicted interacting genes for each prey gene. Genes with a STRING score of 0.95 or greater are displayed.

**Figure S2. Proximity labelling indicates Bcl-2 has a wider subcellular distribution than Bcl-XL.**

**A.** Schematic representation of the BirA*-Bcl-XL and BirA*-Bcl-2 BioID constructs used for experiments in Fig2.

**B.** MCF10A cell lysates from stable lines expressing Venus-BirA*, BirA*-Bcl-XL or BirA*-Bcl-2 immunoblotted with anti Myc confirms equivalent expression levels of each bait. Anti-vinculin was used as a loading control.

**C.** Immunofluorescence imaging of the MCF10A lines in FigS2B. Fixed cells were immunostained with anti-mtHsp70 and anti-Myc-Tag, as indicated.

**D.** Venn diagram comparing the significant hits (B-FDR > 0.05) in the Bcl-XL and Bcl-2 datasets. Prey proteins which are significant in both datasets are identified by the gene name.

**E.** Reactome analysis of significant preys identified in the BirA*-Bcl-2 and BirA*-Bcl-XL BioID analysis.

**F.** Analysis of the biological processes associated with the significantly enriched preys in the BirA*-Bcl-2 and BirA*-Bcl-XL BioID analysis.

**Figure S3. Bax retrotranslocation dependent mitochondrial interaction.**

**A.** Schematic representation of the BirA*-Bax BioID constructs used for experiments in Fig2.

**B.** Isolation of biotin labelled proteins from MCF10A cells stably expressing Venus-BirA*, BirA*-Bcl-XL or BirA*-Bcl-2. Biotin labelled cell lysates from MCF10A cells expressing BirA*-Bax WT, BirA*-BaxS184V and Venus-BirA* were processed on streptavidin-agarose. The input lysate (IN), unbound flow through (FT) and bound and eluted (EL) proteins were separated by SDS-PAGE and blotted with anit-vinculin and biotin, as indicated.

**Figure S4. Generation of Venus-BirA* stable MCF10A cells.**

**A.** Immunofluorescence staining of MCF10A cells stably expressing Venus-BirA* and BadER, and treated with DMSO, 10µM 4-OHT or 5µM ABT-737. In the left panel, cells were co-immunostained with anti-mtHSP70 and anti-myc tag, along with DAPI. Right panel - cells were co-immunostained for the ant-oestrogen receptor (ER) tag to show the BadER fusion protein, alongside anti-mtHSP70 and DAPI. Scale bar = 10µm.

**Figure S5. The BirA*-Bcl-XL proximity interactome shows differential changes when cells are primed by either Bad or the Bad BH3-mimetic, ABT-737.**

**A.** Venn diagram summarising the overlap of significant BioID preys per condition from the BioID data in Fig5A.

**B.** Mapping of the Bcl-xL STRING-predicted interactions (dotted lines) alongside identified BioID hits from Fig5B.

**C)** Mapping of the STRING-predicted interactions of the prey proteins (dotted lines) alongside the identified BioID hits from Fig5B.

**Figure S6. The BirA*-Bcl-2 proximity interactome shows differential changes when cells are primed by either Bad or the Bad BH3-mimetic, ABT-737.**

**A.** Schematic detailing the localisation of Bcl-2 with the different priming treatments and the expression constructs for BirA*-Bcl-2 and BadER.

**B.** Validation of stable MCF10A cells expressing BadER/BirA*-Bcl-2 and BadER/Venus-BirA*. Left panels: Immunofluorescence imaging of both lines with anti-mtHsp70 and anti-Myc-tag, following treatment with DMSO, 10µM 4-OHT or 5µM ABT-737. Scale bar is 10µm. Top right panel: mmunoblots of whole cell lysates of each stable line, using anti-vinculin, anti-Myc-tag, and anti-ER, show similar expression of bait protein and BadER. Bottom right panel: To validate function of BirA*-Bcl-2, MCF10A cells stably expressing BadER/BirA*-Bcl-2 were treated with DMSO, 10μM 4-OHT or 5μM ABT-737 with and without 800μM etoposide. Cells were cultured with FITC-Annexin V and PI and images were taken every hour for 18 hours. The number of Annexin V positive cells counted as a percentage of the total initial cell population. Mean percentages of apoptosis over time are plotted from two independent experiments.

**C.** Volcano plots showing Significance Analysis of Interactome (SAINT) of BioID preys enriched in MCF10A cells expressing BirA*-Bcl-2 relative to Venus-BirA* expressing cells in each of three conditions: cell treated with DMSO; cell treated with 4-OHT to activate BadER; and cells treated with ABT-737. Significant preys (B-FDR > 0,05) indicated in each condition.

**D.** Significant BioID preys for each condition and their overlap between DMSO, 4-OHT and ABT-737 conditions.

**E.** Heatmap showing the relative enrichment of prey proteins across the DMSO, 4-OHT and ABT-737 conditions from Fig.S6C. Each prey is included where it reached the B-FDR > 0.05 in a least one of the conditions in FigS6C.

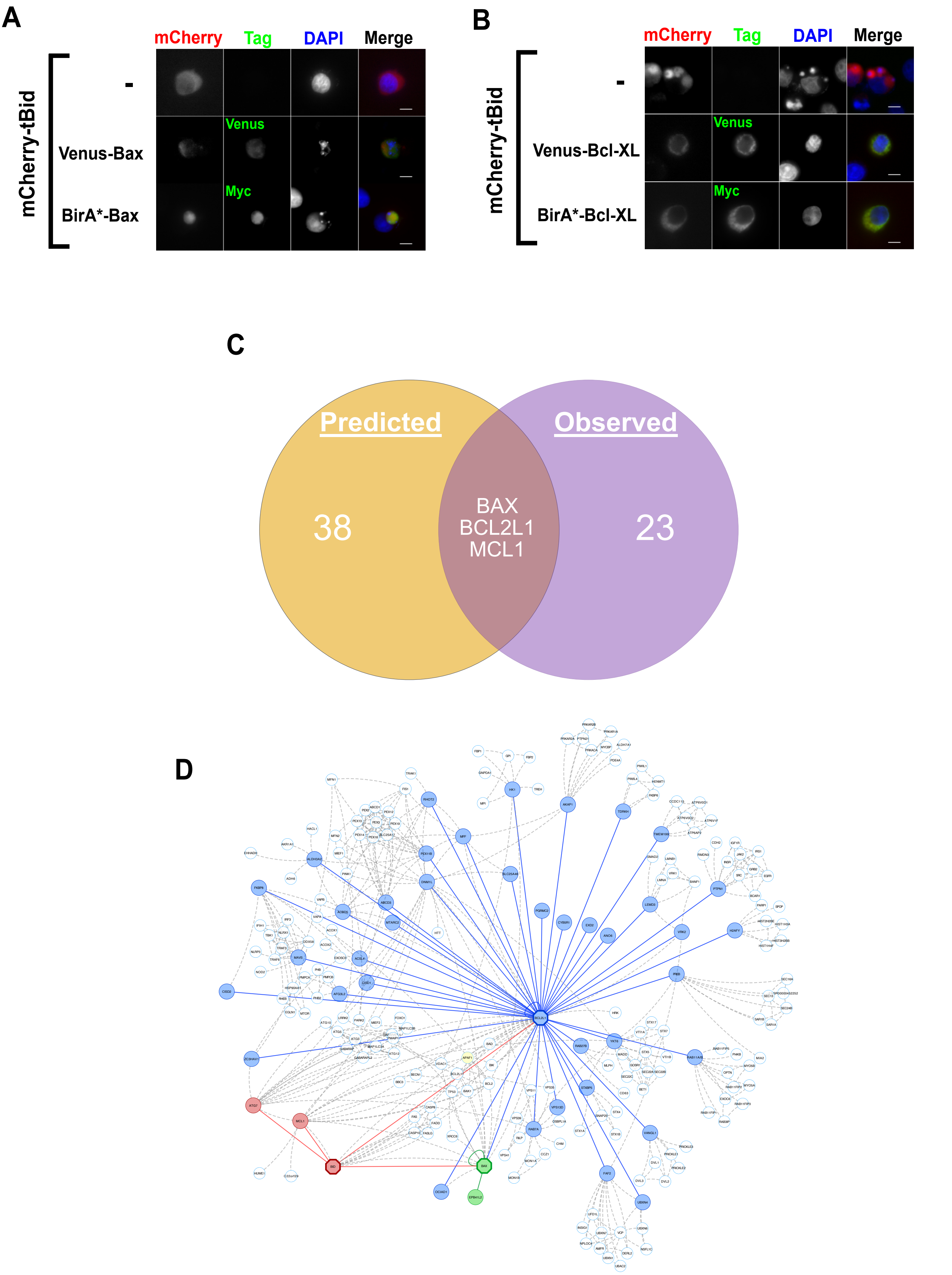
A
mCherry
Tag
DAPI
Merge
mCherry-tBid
-
Venus-Bax
BirA*-Bax
Venus
Myc
B
mCherry
Tag
DAPI
Merge
mCherry-tBid
-
Venus-Bcl-XL
BirA*-Bcl-XL
Venus
Myc
C
Predicted
Observed
38
BAX
BCL2L1
MCL1
23
D

Figure S1

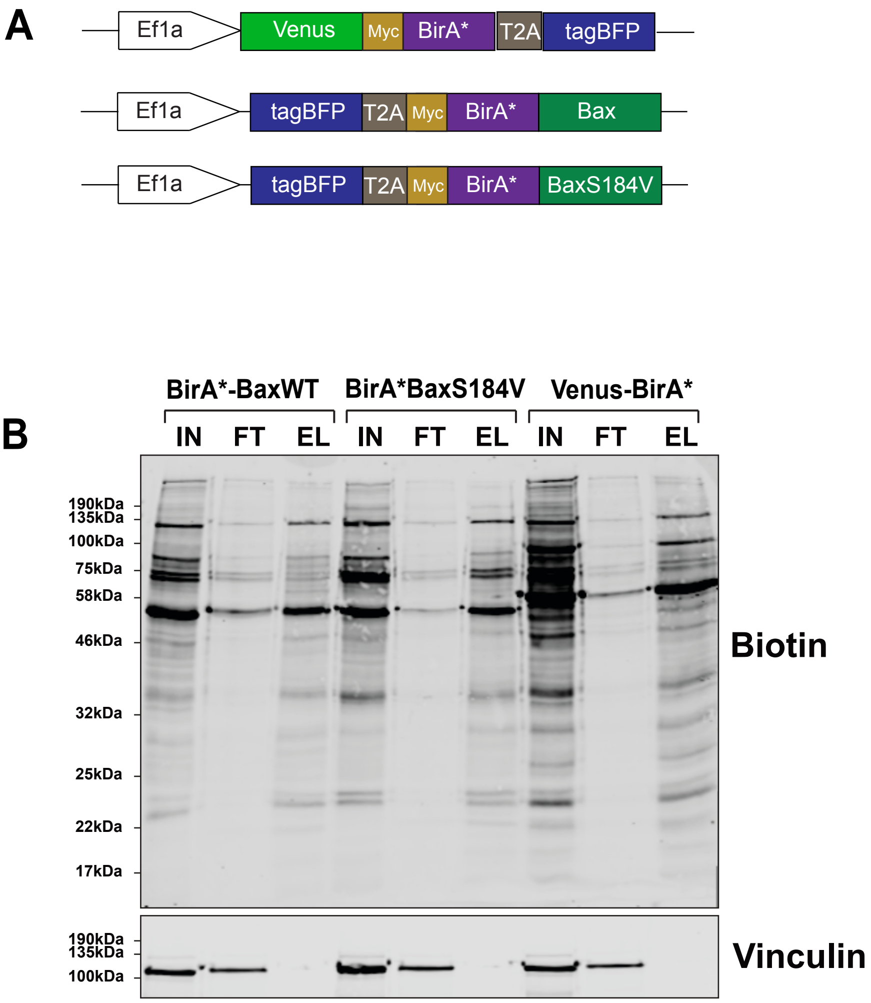
A
Ef1a
Venus
Myc
BirA*
T2A
tagBFP
Bax
BaxS184V
B
BirA*-BaxWT
BirA*BaxS184V
Venus-BirA*
IN
FT
EL
190kDa
135kDa
100kDa
75kDa
58kDa
46kDa
32kDa
25kDa
22kDa
17kDa
Biotin
Vinculin

Figure S2

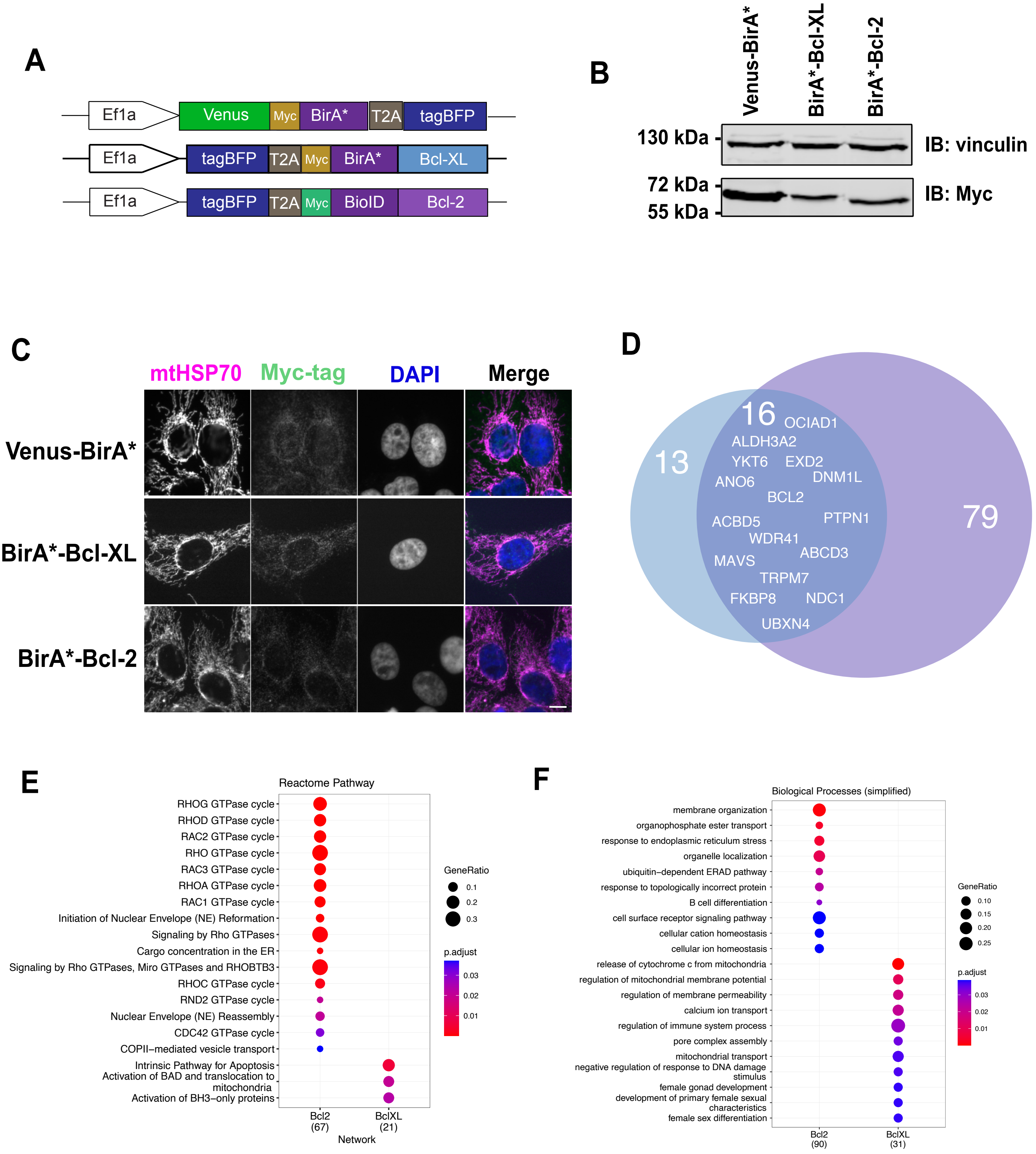
A
Ef1a
Venus
Myc
BirA*
T2A
tagBFP
Ef1a
tagBFP
T2A
Myc
BirA*
Bcl-XL
Ef1a
tagBFP
T2A
Myc
BioID
Bcl-2
B
Venus-BirA*
BirA*-Bcl-XL
BirA*-Bcl-2
130 kDa
IB: vinculin
72 kDa
55 kDa
IB: Myc
C
mtHSP70
Myc-tag
DAPI
Merge
Venus-BirA*
BirA*-Bcl-XL
BirA*-Bcl-2
D
13
16
79
OCIAD1
ALDH3A2
YKT6
EXD2
DNM1L
ANO6
BCL2
ACBD5
PTPN1
WDR41
ABCD3
MAVS
TRPM7
FKBP8
NDC1
UBXN4
E
Reactome Pathway
RHOG GTPase cycle
RHOD GTPase cycle
RAC2 GTPase cycle
RHO GTPase cycle
RAC3 GTPase cycle
RHOA GTPase cycle
RAC1 GTPase cycle
Initiation of Nuclear Envelope (NE) Reformation
Signaling by Rho GTPases
Cargo concentration in the ER
Signaling by Rho GTPases, Miro GTPases and RHOBTB3
RHOC GTPase cycle
RND2 GTPase cycle
Nuclear Envelope (NE) Reassembly
CDC42 GTPase cycle
COPII−mediated vesicle transport
Intrinsic Pathway for Apoptosis
Activation of BAD and translocation to mitochondria
Activation of BH3−only proteins
Bcl2 (67)
BclXL (21)
Network
GeneRatio
0.1
0.2
0.3
p.adjust
0.03
0.02
0.01
F
Biological Processes (simplified)
membrane organization
organophosphate ester transport
response to endoplasmic reticulum stress
organelle localization
ubiquitin−dependent ERAD pathway
response to topologically incorrect protein
B cell differentiation
cell surface receptor signaling pathway
cellular cation homeostasis
cellular ion homeostasis
release of cytochrome c from mitochondria
regulation of mitochondrial membrane potential
regulation of membrane permeability
calcium ion transport
regulation of immune system process
pore complex assembly
mitochondrial transport
negative regulation of response to DNA damage stimulus
female gonad development
development of primary female sexual characteristics
female sex differentiation
Bcl2 (90)
BclXL (31)
GeneRatio
0.10
0.15
0.20
0.25
p.adjust
0.03
0.02
0.01

**Figure S3**

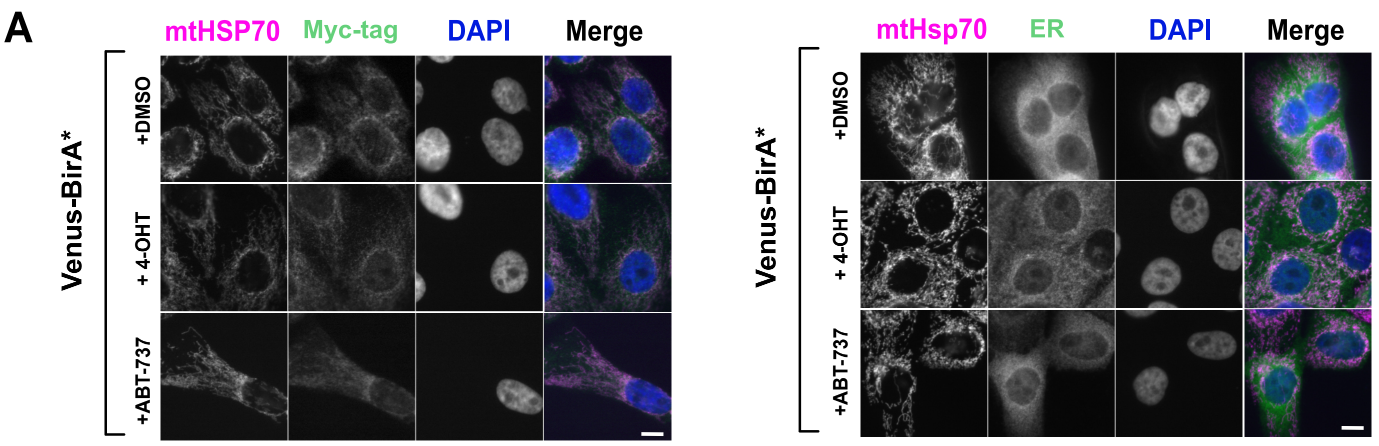
A
mtHSP70
Myc-tag
DAPI
Merge
Venus-BirA*
+DMSO
+ 4-OHT
+ABT-737
mtHsp70
ER
DAPI
Merge
Venus-BirA*
+DMSO
+ 4-OHT
+ABT-737

Figure S4

A

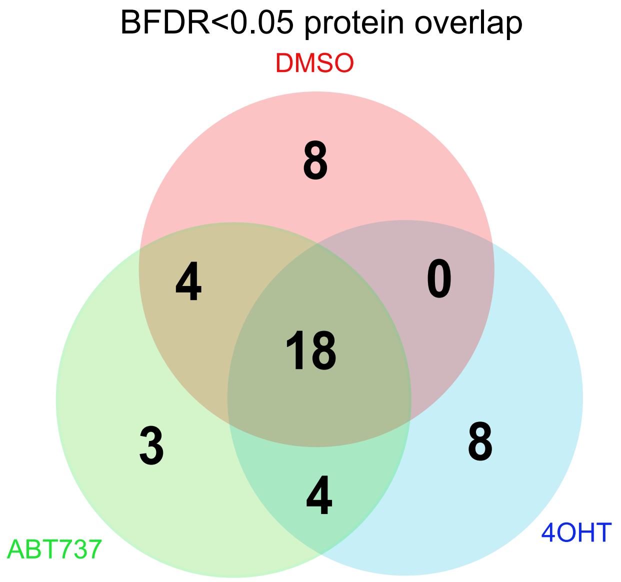


B

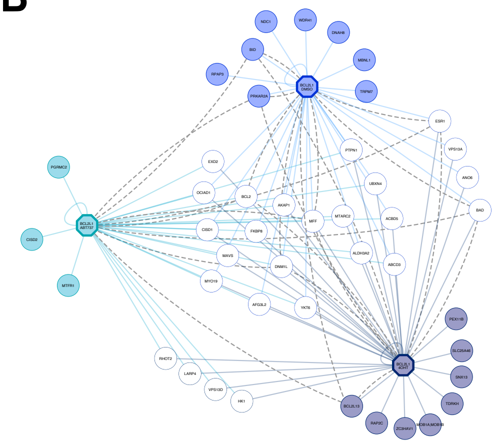

XL STRING predicted connections mapped

C

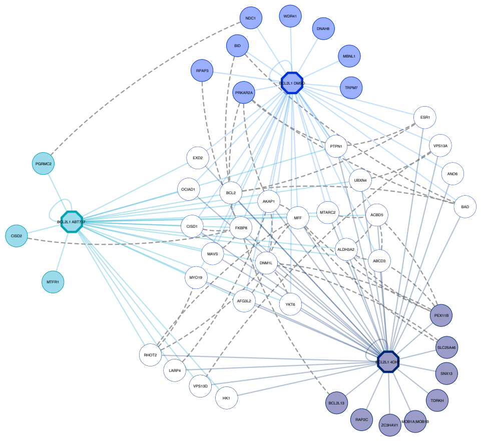

Prey STRING predicted connections mapped

Figure S5

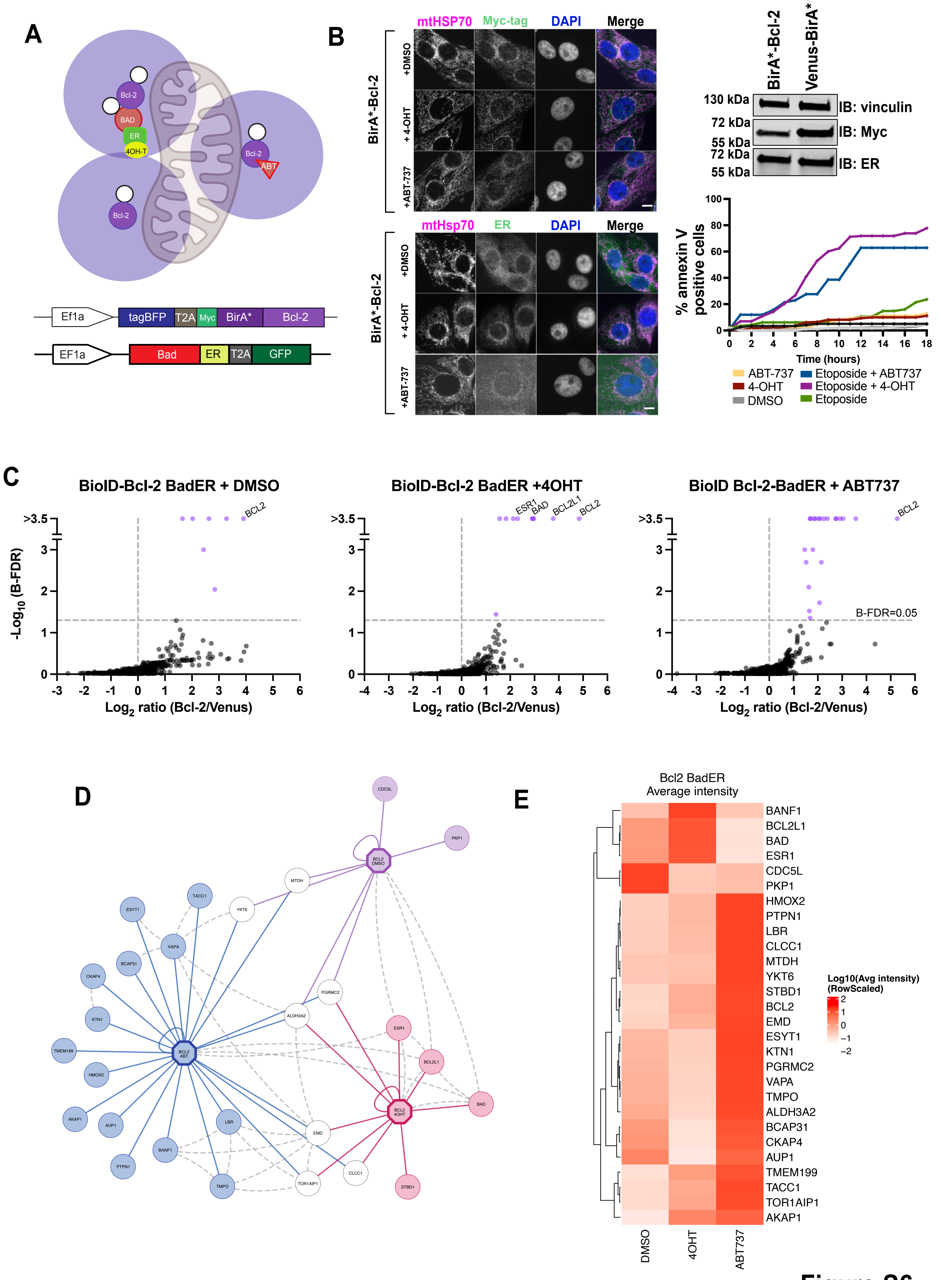

A
Bcl-2
BAD
ER
4OH-T
Bcl-2
ABT
Bcl-2
Ef1a
tagBFP
T2A
Myc
BirA*
Bcl-2
EF1a
Bad
ER
T2A
GFP
B
mtHSP70
Myc-tag
DAPI
Merge
BirA*-Bcl-2
+DMSO
+ 4-OHT
+ABT-737
mtHsp70
ER
DAPI
Merge
BirA*-Bcl-2
BirA*-Bcl-2
Venus-BirA*
130 kDa
IB: vinculin
72 kDa
55 kDa
IB: Myc
72 kDa
55 kDa
IB: ER
% annexin V positive cells
Time (hours)
ABT-737
4-OHT
DMSO
Etoposide + ABT737
Etoposide + 4-OHT
Etoposide
C
BioID-Bcl-2 BadER + DMSO
BioID-Bcl-2 BadER +4OHT
BioID Bcl-2-BadER + ABT737
-Log10 (B-FDR)
Log2 ratio (Bcl-2/Venus)
BCL2
ESR1
BAD
BCL2L1
B-FDR=0.05
D
E
Bcl2 BadER
Average intensity
BANF1
BCL2L1
BAD
ESR1
CDC5L
PKP1
HMOX2
PTPN1
LBR
CLCC1
MTDH
YKT6
STBD1
BCL2
EMD
ESYT1
KTN1
PGRMC2
VAPA
TMPO
ALDH3A2
BCAP31
CKAP4
AUP1
TMEM199
TACC1
TOR1AIP1
AKAP1
Log10(Avg intensity)
(RowScaled)
DMSO
4OHT
ABT737


### Figure S6
